# Object–Action Congruency Shapes Movement Smoothness and Action Preferences in Reachable Space

**DOI:** 10.64898/2026.11.27.769690

**Authors:** Riccardo Moretti, Manuel A. Hernández Alonso, Giovanni Pezzulo, Ignasi Cos

## Abstract

Object affordances can facilitate action execution when the motor response is compatible with the object’s functional properties, but it remains unclear whether these effects extend to naturalistic interactions with physical objects and whether they reflect movement costs. We investigated object– action congruency during simulated functional interactions with a physical mug, its modulation by reachability, and its relationship with movement smoothness and action preferences. Participants performed goal-directed movements to simulate either drinking from a mug or putting it away, with the mug placed horizontally at three reachable distances or at an unreachable location, with its aperture oriented to the left or right. Reaction times and arm kinematics were recorded, and movement smoothness was quantified using log dimensionless jerk. Participants subsequently completed a two-alternative forced-choice task to assess their preferences for previously experienced action– orientation combinations. Congruency effects emerged across multiple kinematic measures. Congruent actions were shorter in movement duration and elicited smoother trajectories than incongruent actions for both goals. Reaction times were shorter for congruent than incongruent stimuli during the Put Away goal, whereas peak velocity was higher for congruent stimuli during the Drink goal; these effects were restricted to reachable distances. In the preference task, participants favoured congruent action–orientation combinations, and choices were predicted by LDJ, indicating a preference for smoother, less costly movements. These findings demonstrate that object–action congruency shapes the execution of naturalistic goal-directed movements and action preferences, with reachability modulating selected kinematic indices. Movement smoothness provides a complementary quantitative measure of congruency, linking motor execution to action selection.

## Introduction

The objects in an environment may offer opportunities for action – or *affordances* (Gibson, 1977). Several studies have shown that object affordances influence behaviour even when irrelevant for the task at hand. For example, a stimulus–response compatibility effect between an object affording a specific grip and its grip response has been reported, mapped to an auditory stimulus (Ellis & Tucker, 2000). In brief, reaction times were reduced whenever the object and motor response associated with the auditory stimulus share a compatible grip. This has been interpreted as reflecting the activation of a motor component—referred to as a micro-affordance—coupled with the object visual representation (Tucker & Ellis, 1998, 2001).

Beyond vision, motor activation is also related to the perception of semantic properties of objects and other cognitive processes, including their identity and function (Creem & Proffitt, 2001; Rounis et al., 2017; Zhang et al., 2021). Some evidence suggests that object perception evokes the canonical object representation (i.e., the one by which the object has been commonly experienced), retrieved from long-term memory, rather than that of the current state of the object. This supports the underlying principle that motor representations are grounded in the distal (i.e., final) state of the action required to render the object functional, as described by the notion of F-commensurability (Masson et al., 2011). Specifically, a grasp has been defined as F-commensurate “when that action results in an outcome that preserves the typical function of the object”. Additional evidence shows that canonical representations are activated regardless of the way the object is oriented (Bub et al., 2015). While these studies included some characterization of movement kinematics, an explicit assessment of movement efficiency (possibly playing a role in the cost-benefit analysis of action) has been overlooked.

Previous research has provided empirical support for the idea that reachable space constitutes an action space (Brozzoli et al., 2013; Bufacchi & Iannetti, 2018; Rizzolatti et al., 1997; Serino, 2026), in which objects may be represented in terms of the actions they afford. Critically, the activation of these action-related representations is not unconditional but appears to abide by adaptive principles. In particular, object reachability plays a key role in determining whether motor information is elicited by visual stimuli. Compatibility assessments between an object and a potential function or manipulation verb are faster when the object is within the reachable space (Costantini et al., 2011). This finding suggests that micro-affordances are conditionally activated, based on the possibility of immediate interaction. Importantly, no such difference was observed for observation verbs, ruling out a purely distance-based principle, and highlighting the relevance of contribution of the action per se. A similar pattern was reported when participants pantomimed reaching movements toward objects that were reachable/unreachable (Costantini et al., 2010). Unreachability was manipulated either by increasing distance or by placing objects behind a semi-transparent barrier. A congruency effect— reflected in faster responses when the object handle was aligned with the responding (ipsilateral) hand—was observed only in the reachable condition. Recently, such a congruency effect (or stimulus alignment effect) and affordance effect (that is the congruency between the response effector and the object’s part of interaction) have been disentangled and observed in both keypress and reach-and-grasp responses in (Demir et al., 2024). Additionally, a fastened response only inside the reachable space has been observed to objects that had less conflict between functional and structural affordances compared to high conflict ones (e.g., objects that afford a structure/shape-related grasp that is different from the one needed to use the object, for instance a calculator) (Kalénine et al., 2016).

However, most of the past experiments have observed congruency effects on simple motor acts (Cavallo et al., 2016) that at most mimicked a few features of a naturalistic goal-directed action in combination with either common daily life objects or towards virtual objects or simple objects with no specific function. The question that remains is whether interacting with a physical object emulating a naturalistic goal-directed action, leads to the same pattern of results. In this regard, (Bub et al., 2021) showed that congruency effects can depend on the type of response required: using identical stimuli, they observed a reverse alignment effect when participants responded by pressing keys, but a positive alignment effect when they performed actual reach-and-grasp actions. This finding suggests that different response modalities may recruit distinct spatial and action-related representations. Accordingly, the first aim of the present study was to assess if the congruency effect is observed when participants interact with a common physical object (e.g., a mug) mimicking a naturalistic goal-directed action. Secondly, we aimed at uncovering whether said congruency effect is modulated by distance and the possibility to reach or not the object. An additional gap in the literature we wanted to address is the absence of a measurable event that allows to characterize quantitatively the concept of congruency beyond the qualitative definitions. In order to achieve this aim, we enriched the classical analysis of reaction times with the inclusion of a set of kinematic variables characterizing the movement kinematics during the task execution. The reason why a kinematic assessment might be foundational to the concept of congruency in object interaction follows from the intuitive understanding of a congruent stimulus as being associated to an easier and more comfortable interaction with the object as compared with an incongruent one. Hence, we propose that a reasonable way to quantify the congruency may be by using a measure of physical effort associated to said object interaction. To approximate that measure we computed the log-dimensionless jerk of the movement trajectory involved in the interaction as a proxy of physical effort. Lastly, we aimed at assessing to what extent participants’ movement preferences were sensitive to the proposed physical effort characterization.

To address these questions, we instructed the participants to interact with a mug placed horizontally on a table, with its aperture oriented towards the right or the left, and at one of three reachable positions or at an unreachable one. We cued one of two possible goals - namely, to simulate drinking from the mug or putting away the mug - before showing the mug, that they had to physically execute. During the interaction with the mug, we recorded the arm kinematic and analysed the reaction time and movement smoothness. Subsequently, to assess participants’ sensitivity to the physical effort metric, we administered a two-alternative forced choice task on a separate laptop. Participants were asked to choose the combination of goal and orientation they preferred out of two options (for all possible combinations they physically experienced during the first task).

We expect that if the naturalistic interactions with physical objects exhibit a congruency effect, participants should show lower reaction times, higher peak velocity, and lower movement duration for congruent stimuli as compared with incongruent ones. Secondly, if the congruency effect is restricted to possible interactions in the reachable space, then it should be observed only at reachable distances, but not for the unreachable one. Third, if the congruency effect is linked to physical effort (or its proxy, movement smoothness), then congruent stimuli within reachable space should elicit smoother (lower cost) trajectories. Lastly, if participants are sensitive to the physical effort of executing the object-directed actions of our task (or its proxy, movement smoothness) they should prefer actions associated with smoother trajectories when given the choice.

## Materials & Methods

### Participants

A sample of 15 participants (6 female) of age 23.8 ± 2.96 years was recruited at the Universitat de Barcelona. All participants gave their written consent by signing an Informed consent form. The participants received a 10€ monetary compensation at the end of the experiment. All the procedures were approved by the Ethics Committee at the Universitat de Barcelona, CER072512, Institutional Review Board IRB00003099.

### Experimental protocol

The experimental session began by providing the participant with task instructions, followed by the reading and signing of the informed consent form. Subsequently, the participant was equipped with a set of 7 passive reflective markers for motion capture using double-sided adhesive tape. The selected anatomical landmarks on the right upper limb were the first metacarpal head, the styloid processes of the radius and ulna, the lateral epicondyle, the midpoint of the posterior surface of the humerus, the acromio-clavicular joint, and the spine of the scapula. An OptiTrack motion capture system comprising six infrared cameras was used to record arm kinematics at a sampling frequency of 100Hz.

The participant sat in front of the table (Fig. 1A) where the experiment was conducted, rested their chin on a chin rest, and wore a pair of computer-controlled liquid crystal occluding goggles (PLATO Visual Occlusion Spectacles, Translucent Technologies Inc.). A forward reach with occluded vision was performed to measure the maximum reachable distance, defined as the distance between the marker on the first metacarpal head and that on the acromio-clavicular joint. This measure was used to define four positions: A, B, C, and D, each corresponding to an offset of -40%, -20%, +20%, and +40% from the maximum reachable distance, respectively. For all participants, positions A, B, and C were reachable (the last requiring participants to fully extend their reach to grasp the mug’s handle, though with a normal power grip), while D was not.

**Figure 1.**
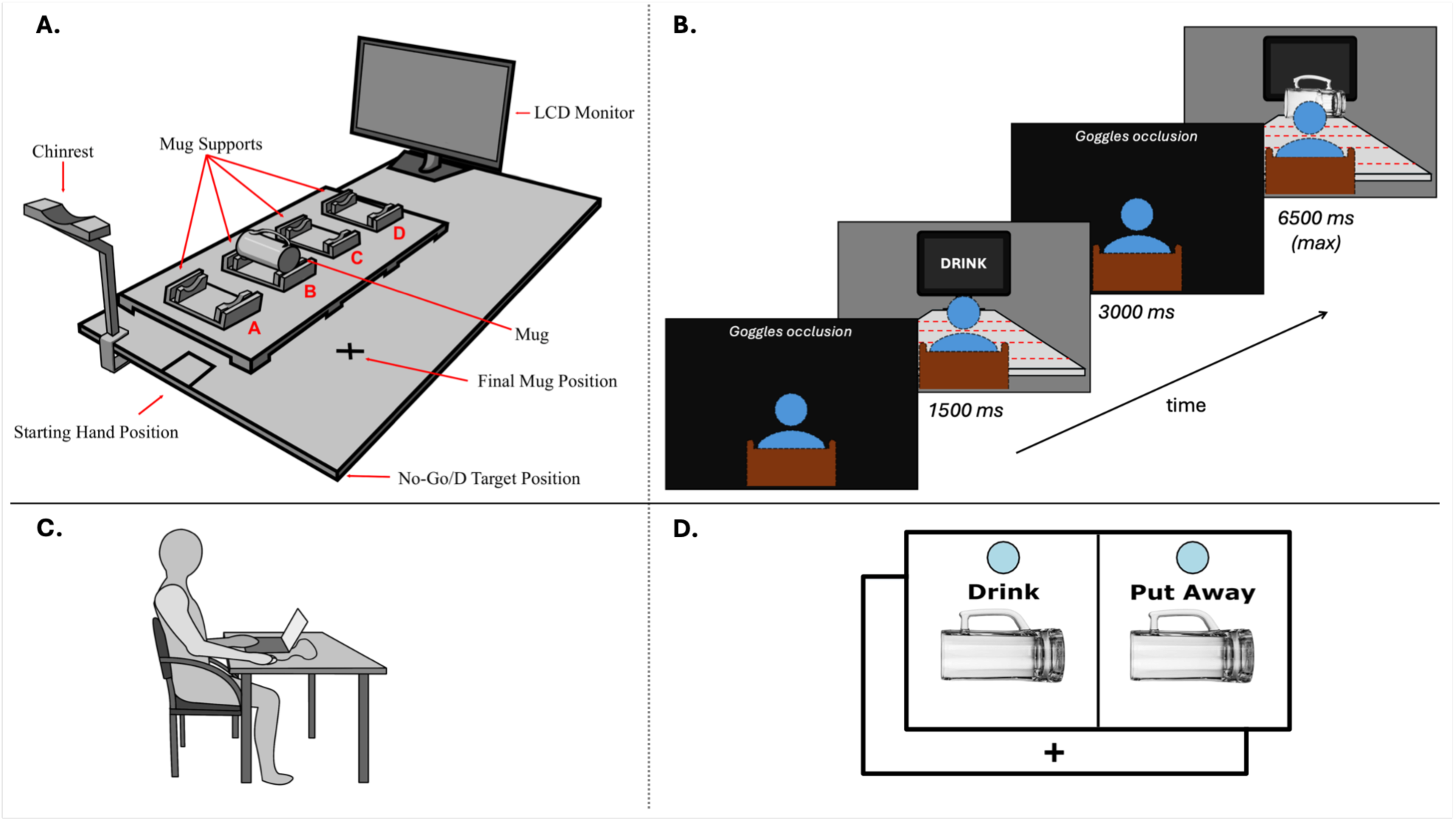
Experimental setup and timeline. **A.** Experimental setup showing the starting position, the four possible distances (i.e., A,B,C, and D) for mug placement (here depicted at position B), the No-Go position and the Final Mug Position. The Participant sat in front of the table with the chin resting on the Chinrest and the right hand resting on the square indicating the Starting Hand Position. **B.** Timeline of a single trial: for 1500ms is shown the cued goal (i.e., Drink or Put Away), next 3000ms of occluded vision are following in order to allow to the experimenter to place the mug in one of the four positions (A,B,C, or D) or on none of them in catch trials (i.e., No-Go condition) and in one of two possible opening orientations (i.e., left or right). After vision is restored, a fast reaction is required to reach and grasp the mug (when reachable) or to move the hand on the No-Go Target position if no mug is on the table or if it is not reachable (position D). The maximum time allowed in this phase is 6500ms, after which a time-out feedback is given to the participant. **C.** Setup for the two alternatives forced choice task (2AFC-task). **D.** Temporal sequence of the 2AFC-task: blank screen with a black cross at the bottom followed by the two options displayed once the participant clicked on the cross. To select one option, it was required to click on the blue circle on top of it. In the example is shown only one possible offer out of six, where both the goal and mug orientation were manipulated.

Before starting, the "action task" was explained to clarify the following sequence of events (Fig. 1B): the trial began with the display, on an LCD monitor positioned at the end of the table, of one of two possible actions to be performed with the mug (placed on one of the four aforementioned positions): to drink from it, or to put it away upside-down on a black cross to the right of the mug support. Vision was then occluded to allow the experimenter to place the mug horizontally — with the handle always oriented upward — on one of the four positions (A, B, C or D), with the mug’s opening facing either left or right. Once the mug was set, vision was restored and participants were required to interact with the mug as quickly as possible, in accordance with the cued action. When participants judged the mug to be unreachable (i.e., in position D) or absent from the table entirely (control No Go condition), the required response was to reach toward the right corner of the table. The combination of 2 actions × 4 distances × 2 orientations yielded 16 conditions. In this set of configurations, half were defined as congruent (Drink-Left and Put Away-Right) and the other half incongruent (Drink-Right and Put Away-Left), the latter requiring a more complex motor pattern to correctly exploit the mug’s function given the cued action.

To clarify the biomechanical rationale underlying the congruent conditions distinction: the starting position consisted of the right hand pronated at a fixed location at the edge of the table in front of the participant (see Starting Hand position in FIG. 1A). The wrist was always aligned with the mug’s handle (both oriented horizontally), but the combination of goal and orientation made pronated grasps biomechanically advantageous for reorienting the object into a functional position. Specifically, the combinations Drink-Left and Put Away-Right were defined as congruent because, once grasped with a pronated grip (consistent with the forearm orientation at the starting position), a 90° clockwise rotation was sufficient to bring the mug into its functional end-state orientation. Conversely, the two remaining combinations, Drink-Right and Put Away-Left, required rotating the forearm into a supinated position prior to grasping, in order for a simple 90° counter-clockwise rotation to restore the object to a functional state; this additional complexity renders them incongruent in the present context in accordance with F-commensurability as defined by (Masson et al., 2011).

In addition, there was the No Go condition × 2 goals, for a total of 18 conditions. The number of trials for each condition was 20, for a total of 360 trials distributed across 5 blocks of 72 trials, with a 2-minute rest between blocks. This phase of the experiment lasted approximately one hour.

Following the action task, the Edinburgh Handedness Inventory (Oldfield, 1971) and a second task, a two-alternative forced choice task (2AFC-task) lasting ten minutes, were administered.

The 2AFC-task was performed on a laptop (FIG. 1C). Each of the two options displayed an image of the mug, always shown with the handle oriented upward and the opening facing either left or right (FIG. 1D). One of the two actions (to drink or to put away) was displayed above the image, such that each option corresponded to a combination of goal and orientation, as in the "action task", with the exception that distance was not manipulated. The total number of available options was 4, yielding 6 possible pairings of two options out of four, each offer repeated 20 times in pseudorandom counterbalanced order. Participants were asked to select the option they would prefer to execute (ideally, the one subjectively perceived as less effortful), by clicking the corresponding button with the mouse.

### Action task

We analysed the marker trajectories by first smoothing them with a Gaussian filter with a sigma of five. Each marker trajectory was recorded as a 3D vector [x, y, z], tracked at 100Hz by six Optitrack cameras (Optitrack, Inc.), and referenced with respect to the shoulder’s marker position (as the origin of movement). The norm vector of each marker was then computed. Velocity, acceleration, and jerk were subsequently derived from the norm by numerical differentiation. The velocity profile was subsequently used to detect movement onset, defined as the point at which velocity exceeded a threshold of 3% of peak absolute velocity. Reaction time (RT) was defined as the time interval between vision restoration and movement onset. Movement duration (MD) was defined as the time interval between RT and the return of the hand to the starting position. Additional kinematic variables computed were: peak velocity (PV), absolute time to peak velocity (aTTPV), peak acceleration (PA), absolute time to peak acceleration (aTTPA), movement trajectory length (ML), and log-dimensionless jerk (LDJ) (Melendez-Calderon et al., 2021), the latter being used to quantify movement smoothness as a proxy for movement cost. The aforementioned variables were computed for each marker, and the statistical comparison between congruent and incongruent trials took into account the full set of markers across the arm to capture the effect of the experimental manipulation on whole-arm kinematics. Prior to hypothesis testing, for each participant and anatomical marker, outliers were removed by excluding trajectories with a movement onset earlier than 150ms to exclude anticipatory responses (similarly to the threshold used in (Contemori et al., 2021)) and values below Q1 − (1.5 × IQR) or above Q3 + (1.5 × IQR) for each variable. IQR stands for the interquartile range and has been computed as the difference between the values corresponding to the third (Q3) and first (Q1) quartiles.

### Statistical Analyses

We assessed the effect of our experimental conditions on the aforementioned variables by fitting Linear Mixed-Effects Models, implemented with the *lme4* (Bates et al., 2015) and *lmerTest* (Kuznetsova et al., 2017) R packages. Models were fit using restricted maximum likelihood (REML) and included all fixed effects (i.e., Distance, Orientation, and Goal) and their two-way and three-way interactions (Equation 1). P-values for fixed effects were derived from t-tests using the Satterthwaite approximation for degrees of freedom. To account for inter-individual variability and to capture group-level effects, a random intercept per subject was included in each model. All models set the reference level at the combination of Distance = A, Orientation = Left, and Goal = Drink, such that fixed-effect coefficients represent the effect of each condition relative to this baseline.

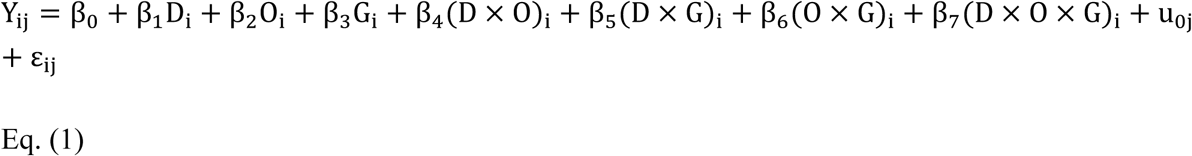

Abbreviations: D, Distance; O, Orientation; G, Goal; Y, response variable (dependent measure); β₀, fixed intercept; β₁–β₃, fixed main effects of Distance, Orientation, and Goal, respectively; β₄–β₆, fixed two-way interaction effects; β₇, fixed three-way interaction effect; u_0j_, random intercept for subject j; ε_ij_, residual error term.

Post-hoc pairwise comparisons of estimated marginal means (Equation 2) were conducted using the *emmeans* R package (Lenth & Piaskowski, 2026) to assess the magnitude of the congruency effect at each distance, for each goal, with Bonferroni correction applied for multiple comparisons. Because the denominator degrees of freedom exceeded the threshold for the t-to-z asymptotic approximation, pairwise contrasts were evaluated using z-tests.

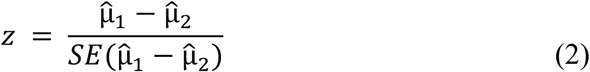

## Results

This section describes the results for the Movement Duration, Reaction Time, Peak Velocity and Log-Dimensionless Jerk. The analysis of the remaining kinematic variables (i.e., Movement Length, aTTPV, PA and aTTPA) can be found in the Supplementary Results. Moreover, the analysis of decisions from the two alternatives forced choice task are presented at the end of this section.

### Movement duration

The LMM on MD (Fig 2A, Table 1) revealed a statistically significant main effect of Distance, whereby movement duration decreased at distance **B** (β = -0.095, *SE* = 0.027, *t*(28439) = -3.511, *p* = 4.0 × 10^-4^, 95% *CI* [-0.148, -0.042]), to a smaller extent at distance **C** (β = -0.054, *SE* = 0.028, *t*(28439) = -1.972, *p* = 0.049, 95% *CI* [-0.108, -3.0 × 10^-4^]) and greatly at distance **D** (β = -1.487, *SE* = 0.027, *t*(28439) = -55.524, *p* = 0.0000, 95% *CI* [-1.54, -1.435]). Distance D required a simple straight movement to the right corner of the table and therefore exhibited a smaller duration than all other conditions.

**Figure 2.**
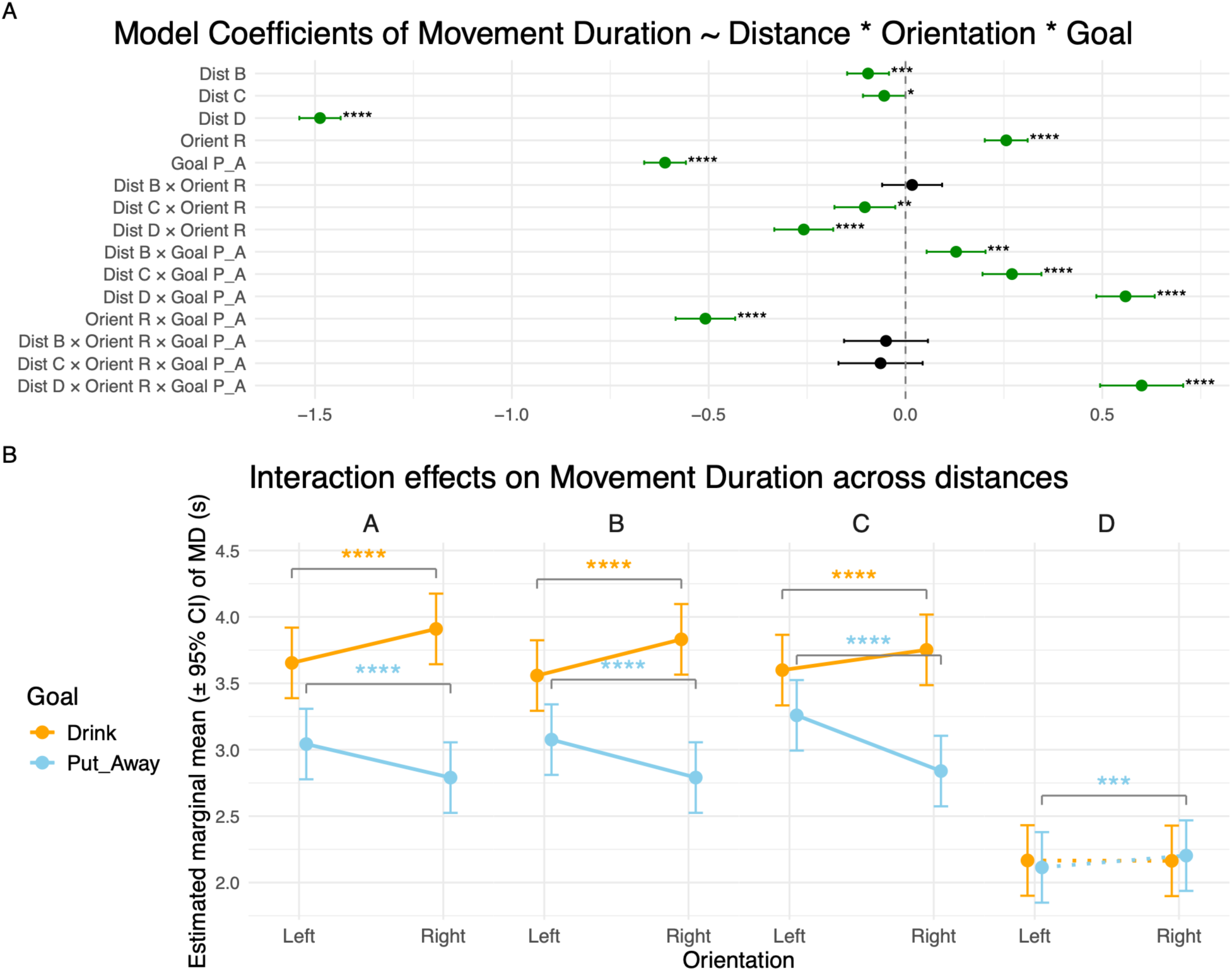
Linear Mixed Model results for Movement Duration. **A)** Forest plot of the LMM coefficients for Movement Duration, modelled as a function of Distance, Orientation, and Goal, including all interaction terms. Each row represents one predictor or interaction term, with the point estimate and 95% confidence interval displayed horizontally. Statistically significant coefficients are shown in green; the non-significant coefficient is shown in black. Asterisks denote the level of significance (* p ≤ 0.05, ** p ≤ 0.01, *** p ≤ 0.001, **** p ≤ 0.0001). **B)** Estimated marginal means (± 95% CI) of Movement Duration as a function of Orientation (Left, Right) at each distance (A, B, C, D), separately for each Goal (Drink: orange; Put Away: sky blue). Brackets with asterisks indicate planned pairwise comparisons between Left and Right orientations within each Distance × Goal combination, with the same significance coding as in panel A. Distance D is shown with dashed lines as it corresponds to a movement directed toward the right corner of the table rather than toward the mug. Note: the **congruent** combinations of Goal and Orientation are represented by Drink-Left and Put Away-Right, the **incongruent** combinations of Goal and Orientation are represented by Drink-Right and Put Away-Left.

**Table 1.** Linear Mixed Model coefficients for Movement Duration. Fixed-effect estimates from the LMM with Movement Duration as the dependent variable, modelled as a function of Distance, Orientation, Goal, and all interaction terms. For each predictor, the table reports the coefficient estimate, standard error (SE), degrees of freedom (df), t-value, p-value, and the lower and upper bounds of the 95% confidence interval (CI). Subject ID was included as a random effect. The model intercept corresponds to the combination of Distance = A, Orientation = Left, and Goal = Drink. Distance, Orientation, and Goal are abbreviated as Dist, Orient, and Goal, respectively.

| Model — MD ~ Distance × Orientation × Goal |  |  |  |  |  |  |  |
| Fixed Effect | Estimate | SE | df | t | p | Lower 95% CI | Upper 95% CI |
|---|---|---|---|---|---|---|---|
| (Intercept) | 3.6538 | 0.1355 | 14,6 | 26.961 | **7.83e-14** | 3.3642 | 3.9434 |
| Dist B | -0.0952 | 0.0271 | 28439 | -3.511 | **0.0004** | -0.1483 | -0.0420 |
| Dist C | -0.0542 | 0.0275 | 28439 | -1.972 | **0.0486** | -0.1081 | -0.0003 |
| Dist D | -1.4874 | 0.0268 | 28439 | -55.524 | **0.0000** | -1.5400 | -1.4349 |
| Orient R | 0.2558 | 0.0278 | 28439 | 9.196 | **3.95e-20** | 0.2013 | 0.3103 |
| Goal P_A | -0.6109 | 0.0270 | 28439 | -22.606 | **3.72e-112** | -0.6639 | -0.5580 |
| Dist B × Orient R | 0.0167 | 0.0389 | 28439 | 0.430 | 0.6669 | -0.0595 | 0.0930 |
| Dist C × Orient R | -0.1033 | 0.0394 | 28439 | -2.623 | **0.0087** | -0.1806 | -0.0261 |
| Dist D × Orient R | -0.2585 | 0.0382 | 28439 | -6.767 | **1.34e-11** | -0.3334 | -0.1836 |
| Dist B × Goal P_A | 0.1284 | 0.0382 | 28439 | 3.364 | **0.0008** | 0.0536 | 0.2032 |
| Dist C × Goal P_A | 0.2704 | 0.0382 | 28439 | 7.082 | **1.46e-12** | 0.1956 | 0.3452 |
| Dist D × Goal P_A | 0.5589 | 0.0378 | 28439 | 14.772 | **3.39e-49** | 0.4847 | 0.6330 |
| Orient R × Goal P_A | -0.5084 | 0.0386 | 28439 | -13.172 | **1.66e-39** | -0.5841 | -0.4328 |
| Dist B × Orient R × Goal P_A | -0.0496 | 0.0544 | 28439 | -0.912 | 0.3617 | -0.1562 | 0.0570 |
| Dist C × Orient R × Goal P_A | -0.0634 | 0.0546 | 28439 | -1.163 | 0.2450 | -0.1704 | 0.0435 |
| Dist D × Orient R × Goal P_A | 0.5998 | 0.0538 | 28439 | 11.139 | **9.32e-29** | 0.4943 | 0.7054 |

The main effects of Orientation Right (β = 0.256, *SE* = 0.028, *t*(28439) = 9.196, *p* = 3.95 × 10^-20^, 95% *CI* [0.201, 0.31]) and Goal Put Away (β = -0.611, *SE* = 0.027, *t*(28439) = -22.606, *p* = 3.72 × 10^-112^, 95% *CI* [-0.664, -0.558]) were also statistically significant, consistently with movement duration lasting longer for movements involving a right-oriented mug compared with a left-oriented mug. Movement duration also differed between goals, being smaller for the Put Away goal than for the Drink goal.

Interestingly we observed a two-way interaction between Orientation Right and Goal Put Away (that is a congruent combination), yielding a shorter movement duration (β = -0.508, *SE* = 0.039, *t*(28439) = -13.172, *p* = 1.66 × 10^-39^, 95% *CI* [-0.584, -0.433]).

The pairwise comparison (Fig.2B, Table 2) was significant at every reachable distance for both goals, supporting the congruency effect. The differences between congruency conditions for the goal Drink (Δ = congruent - incongruent) resulted in: **A** (Δ = -0.256, *SE* = 0.029, *z* = -9.196, *p_Bonferroni_* = 3.7 × 10^-20^, 95% *CI* [-0.31, -0.201]), **B** (Δ = -0.273, *SE* = 0. 027, *z* = -10.026, *p_Bonferroni_* = 1.17 × 10^-23^, 95% *CI* [-0.326, -0.219]) and **C** (Δ = -0.152, *SE* = 0.028, *z* = -5.461, *p_Bonferroni_* = 4.73 × 10^-8^, 95% *CI* [-0.207, -0.098]).

**Table 2.** Pairwise comparisons of Movement Duration between Left and Right orientations. Estimated marginal mean differences (Δ = Left − Right) between orientation conditions at each Distance × Goal combination (2 Goals × 4 Distances), derived from the LMM using the emmeans package. P-values are Bonferroni-corrected for multiple comparisons. Note: For the Drink goal, Left − Right corresponds to Δ = congruent − incongruent; for the Put Away goal, Left − Right corresponds to Δ = incongruent − congruent.

| **Pairwise Comparisons — MD** |  |  |  |  |  |  |  |  |
|---|---|---|---|---|---|---|---|---|
| **Contrast** | **Goal** | **Distance** | **Estimate** | **SE** | **z** | **p** | **Lower 95% CI** | **Upper 95% CI** |
| Left - Right | Drink | A | -0.2558 | 0.0278 | -9.196 | **3.70e-20** | -0.3103 | -0.2013 |
| Left - Right | Drink | B | -0.2725 | 0.0272 | -10.026 | **1.17e-23** | -0.3258 | -0.2192 |
| Left - Right | Drink | C | -0.1524 | 0.0279 | -5.461 | **4.73e-08** | -0.2071 | -0.0977 |
| Left - Right | Drink | D | 0.0027 | 0.0262 | 0.105 | 0.9166 | -0.0486 | 0.0541 |
| Left - Right | Put_Away | A | 0.2526 | 0.0267 | 9.449 | **3.41e-21** | 0.2002 | 0.3050 |
| Left - Right | Put_Away | B | 0.2855 | 0.0269 | 10.597 | **3.08e-26** | 0.2327 | 0.3383 |
| Left - Right | Put_Away | C | 0.4194 | 0.0266 | 15.776 | **4.54e-56** | 0.3673 | 0.4715 |
| Left - Right | Put_Away | D | -0.0886 | 0.0269 | -3.300 | **0.0010** | -0.1413 | -0.0360 |

The differences between congruency conditions for the goal Put_Away (Δ = incongruent - congruent) were partially consistent with congruency effect predictions. For the goal Put_Away at the distances **A** (Δ = 0.253, *SE* = 0.027, *z* = 9.449, *p_Bonferroni_* = 3.41 × 10^-21^, 95% *CI* [0.2, 0.305]), **B** (Δ = 0.286, *SE* = 0.027, *z* = 10.597, *p_Bonferroni_* = 3.08 × 10^-26^, 95% *CI* [0.233, 0.338]) and **C** (Δ = 0.419, *SE* = 0.027, *z* = 15.776, *p_Bonferroni_* = = 4.54 × 10^-56^, 95% *CI* [0.367, 0.472]). Unexpectedly we observed a statistically significant difference for the goal Put Away at distance **D** (Δ = -0.089, *SE* = 0.027, *z* = -3.3, *p_Bonferroni_* = 0.001, 95% *CI* [-0.141, 0.036]).

In summary, the congruent conditions were associated with a shorter movement duration.

### Reaction time

We assessed the dependence of RT on Distance, Orientation and Goal by means of a LMM (Fig.3A, Table 3). This revealed significant main effects of Distance and Goal. Specifically, RT increased relative to the closest distance (**A**) at distance **C** (β = 0.052, *SE* = 0.005, *t*(27957) = 9.648, *p* = 5.45 × 10^-22^, 95% *CI* [0.042, 0.063]) and to a greater extent at **D** (β = 0.117, *SE* = 0.005, *t*(27957) = 21.965, *p* = 4.93 × 10^-106^, 95% *CI* [0.107, 0.128]), and decreased at **B** (β = -0.012, *SE* = 0.005, *t*(27957) = - 2.307, *p* = 0.021, 95% *CI* [-0.023, -0.002]). The Put Away goal yielded a longer RT compared to the Drink goal (β = 0.032, *SE* = 0.005, *t*(27957) = 5.895, *p* = 3.78 × 10^-9^, 95% *CI* [0.021, 0.042]). The interaction between Orientation Right and Goal Put away (that represents a congruent combination) resulted in a statistically significant coefficient (β = -0.025, *SE* = 0.008, *t*(27957) = -3.314, *p* = 9.2 × 10^-4^, 95% *CI* [-0.04, -0.01]), denoting shorter RTs for congruent combinations.

**Figure 3.**
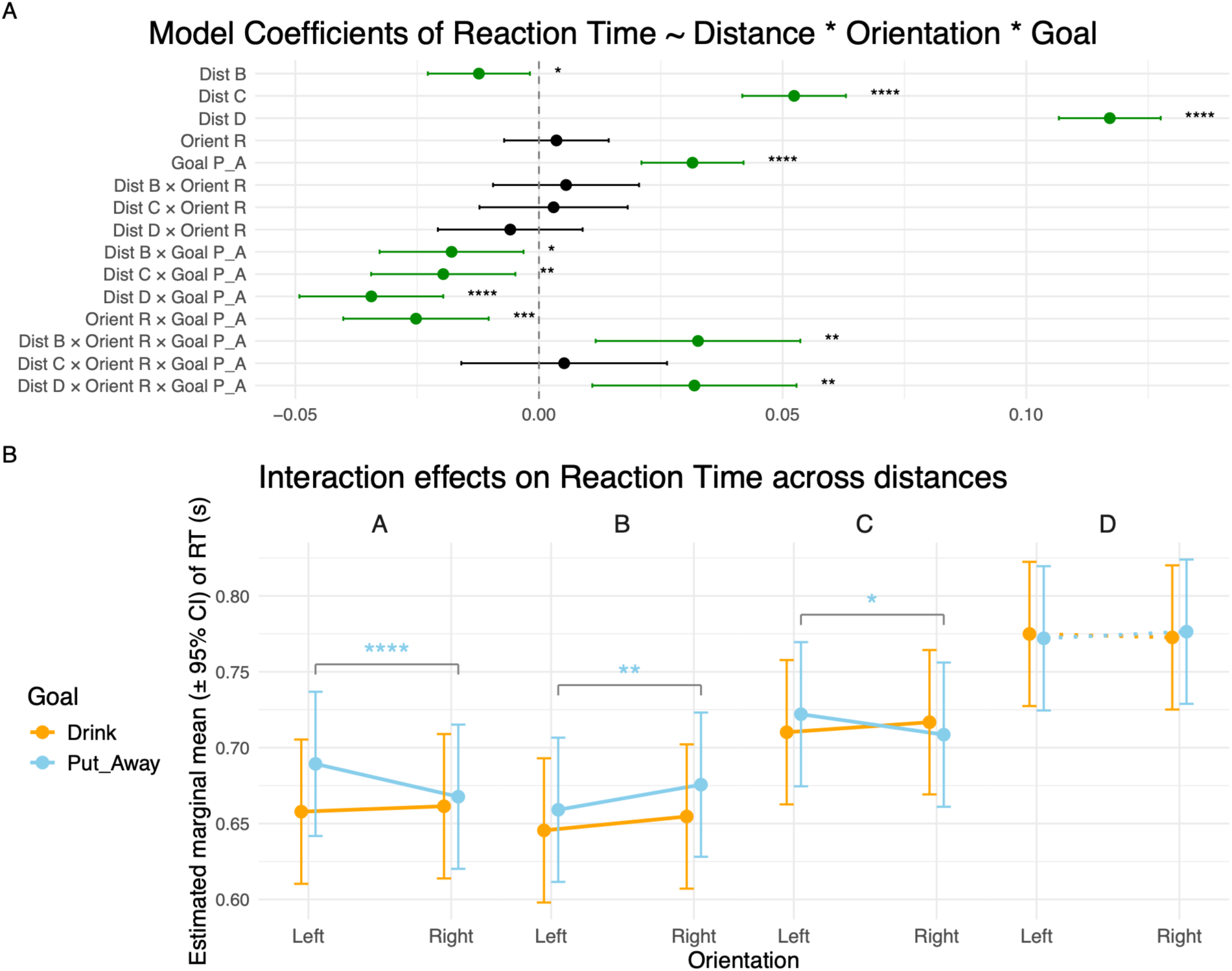
Linear Mixed Model results for Reaction Time (RT). **A.** Forest plot of the LMM coefficients (mean and standard error) for RT, regressed as a function of Distance, Orientation, and Goal, including all interaction terms. Each row represents one predictor or interaction term, with the point estimate and 95% confidence interval displayed horizontally. Statistically significant coefficients are shown in green; the non-significant coefficient is shown in black. Asterisks denote the level of significance (* p ≤ 0.05, ** p ≤ 0.01, *** p ≤ 0.001, **** p ≤ 0.0001). **B.** Estimated marginal means (± 95% CI) of RT as a function of Orientation (Left, Right) at each distance (A, B, C, D), separately for each Goal (Drink: orange; Put Away: sky blue). Asterisks indicate planned pairwise comparisons between Left and Right orientations within each Distance × Goal combination, with the same significance coding as in panel A. Distance D is shown with dashed lines as it corresponds to a movement directed toward the right corner of the table rather than toward the mug. Note: the **congruent** combinations of Goal and Orientation are represented by Drink-Left and Put Away-Right, the **incongruent** combinations of Goal and Orientation are represented by Drink-Right and Put Away-Left.

**Table 3.** Linear Mixed Model coefficients for RT. Fixed-effect estimates from the LMM with Reaction Time as the dependent variable, modelled as a function of Distance, Orientation, Goal, and all interaction terms. For each predictor, the table reports the coefficient estimate, standard error (SE), degrees of freedom (df), t-value, p-value, and the lower and upper bounds of the 95% confidence interval (CI). Subject ID was included as a random effect. The model intercept corresponds to the combination of Distance = A, Orientation = Left, and Goal = Drink. Distance, Orientation, and Goal are abbreviated as Dist, Orient, and Goal, respectively.

| **Model — RT ~ Distance × Orientation × Goal** |  |  |  |  |  |  |  |
| **Fixed Effect** | **Estimate** | **SE** | **df** | **t** | **p** | **Lower 95% CI** | **Upper 95% CI** |
|---|---|---|---|---|---|---|---|
| (Intercept) | 0.6578 | 0.0243 | 14,7 | 27.118 | **6.05e-14** | 0.6060 | 0.7096 |
| Dist B | -0.0123 | 0.0053 | 27957 | -2.307 | **0.0211** | -0.0228 | -0.0019 |
| Dist C | 0.0524 | 0.0054 | 27957 | 9.648 | **5.45e-22** | 0.0417 | 0.0630 |
| Dist D | 0.1171 | 0.0053 | 27957 | 21.965 | **4.93e-106** | 0.1067 | 0.1276 |
| Orient R | 0.0036 | 0.0055 | 27957 | 0.655 | 0.5122 | -0.0071 | 0.0143 |
| Goal P_A | 0.0315 | 0.0053 | 27957 | 5.895 | **3.78e-09** | 0.0210 | 0.0420 |
| Dist B × Orient R | 0.0056 | 0.0076 | 27957 | 0.728 | 0.4667 | -0.0094 | 0.0205 |
| Dist C × Orient R | 0.0030 | 0.0078 | 27957 | 0.387 | 0.6987 | -0.0122 | 0.0182 |
| Dist D × Orient R | -0.0059 | 0.0076 | 27957 | -0.777 | 0.4373 | -0.0207 | 0.0090 |
| Dist B × Goal P_A | -0.0179 | 0.0075 | 27957 | -2.379 | **0.0174** | -0.0327 | -0.0032 |
| Dist C × Goal P_A | -0.0196 | 0.0076 | 27957 | -2.601 | **0.0093** | -0.0344 | -0.0048 |
| Dist D × Goal P_A | -0.0344 | 0.0075 | 27957 | -4.568 | **4.96e-06** | -0.0492 | -0.0196 |
| Orient R × Goal P_A | -0.0252 | 0.0076 | 27957 | -3.314 | **0.0009** | -0.0402 | -0.0103 |
| Dist B × Orient R × Goal P_A | 0.0326 | 0.0107 | 27957 | 3.046 | **0.0023** | 0.0116 | 0.0536 |
| Dist C × Orient R × Goal P_A | 0.0052 | 0.0108 | 27957 | 0.480 | 0.6309 | -0.0159 | 0.0263 |
| Dist D × Orient R × Goal P_A | 0.0319 | 0.0107 | 27957 | 2.984 | **0.0029** | 0.0109 | 0.0529 |

The pairwise comparison (Fig.3B, Table 4) was significant at every reachable distance for the Put Away goal. The differences between congruency conditions for Put_Away (Δ = incongruent - congruent) were partially consistent with prediction. For the goal Put_Away at the distances **A** (Δ = 0.022, *SE* = 0.005, *z* = 4.095, *p_Bonferroni_* = 4.22 × 10^-5^, 95% *CI* [0.011, 0.032]) and **C** (Δ = 0.014, *SE* = 0.005, *z* = 2.559, *p_Bonferroni_* = 0.011, 95% *CI* [0.003, 0.024]), the effect was consistent with our prediction: the congruent condition yielded a faster RT. For distance **B** (Δ = -0.017, *SE* = 0.005, *z* = -3.113, *p_Bonferroni_* = 0.002, 95% *CI* [-0.027, -0.006]), the reverse pattern was observed. At distance **D** (Δ = -0.004, *SE* = 0.005, *z* = -0.814, *p_Bonferroni_* = 0.416, 95% *CI* [-0.015, 0.006]), no effect was observed. For the Drink goal, the expected congruency pattern was observed only qualitatively, with none of the differences reaching statistical significance.

**Table 4.** Pairwise comparisons of Reaction Time between Left and Right orientations. Estimated marginal mean differences (Δ = Left − Right) between orientation conditions at each Distance × Goal combination (2 Goals × 4 Distances), derived from the LMM using the emmeans package. P-values are Bonferroni-corrected for multiple comparisons. Note: For the Drink goal, Left − Right corresponds to Δ = congruent − incongruent; for the Put Away goal, Left − Right corresponds to Δ = incongruent − congruent.

| **Pairwise Comparisons — RT** |  |  |  |  |  |  |  |  |
| **Contrast** | **Goal** | **Distance** | **Estimate** | **SE** | **z** | **p** | **Lower 95% CI** | **Upper 95% CI** |
|---|---|---|---|---|---|---|---|---|
| Left - Right | Drink | A | -0.0036 | 0.0055 | -0.655 | 0.5122 | -0.0143 | 0.0071 |
| Left - Right | Drink | B | -0.0092 | 0.0053 | -1.717 | 0.0860 | -0.0196 | 0.0013 |
| Left - Right | Drink | C | -0.0066 | 0.0055 | -1.199 | 0.2305 | -0.0174 | 0.0042 |
| Left - Right | Drink | D | 0.0023 | 0.0052 | 0.439 | 0.6610 | -0.0080 | 0.0126 |
| Left - Right | Put_Away | A | 0.0216 | 0.0053 | 4.095 | **4.22e-05** | 0.0113 | 0.0320 |
| Left - Right | Put_Away | B | -0.0166 | 0.0053 | -3.113 | **0.0018** | -0.0270 | -0.0061 |
| Left - Right | Put_Away | C | 0.0135 | 0.0053 | 2.559 | **0.0105** | 0.0032 | 0.0238 |
| Left - Right | Put_Away | D | -0.0044 | 0.0054 | -0.814 | 0.4158 | -0.0149 | 0.0062 |

In summary, we observed a congruency effect within the reachable space for the Put Away goal but no effect outside the participants’ reach, supporting the conclusion that the congruency effect is conditional on the perceived possibility of interacting with the object.

### Peak velocity

The LMM on peak velocity (Fig 4A, Table 5) revealed a statistically significant main effect of Distance, whereby PV increased at distances **B** (β = 0.222, *SE* = 0.015, *t*(28359) = 15.101, *p* = 2.53 × 10^-51^, 95% *CI* [0.193, 0.25]), and **C** (β = 0.417, *SE* = 0.015, *t*(28359) = 27.982, *p* = 5.48 × 10^-170^, 95% *CI* [0.388, 0.447]) and decreased at distance **D** (β = -0.041, *SE* = 0.015, *t*(28360) = -2.798, *p* = 0.005, 95% *CI* [-0.069, -0.012]). Distance D required a simple straight movement to the right corner of the table and exhibited a smaller peak velocity than all other distances.

**Figure 4.**
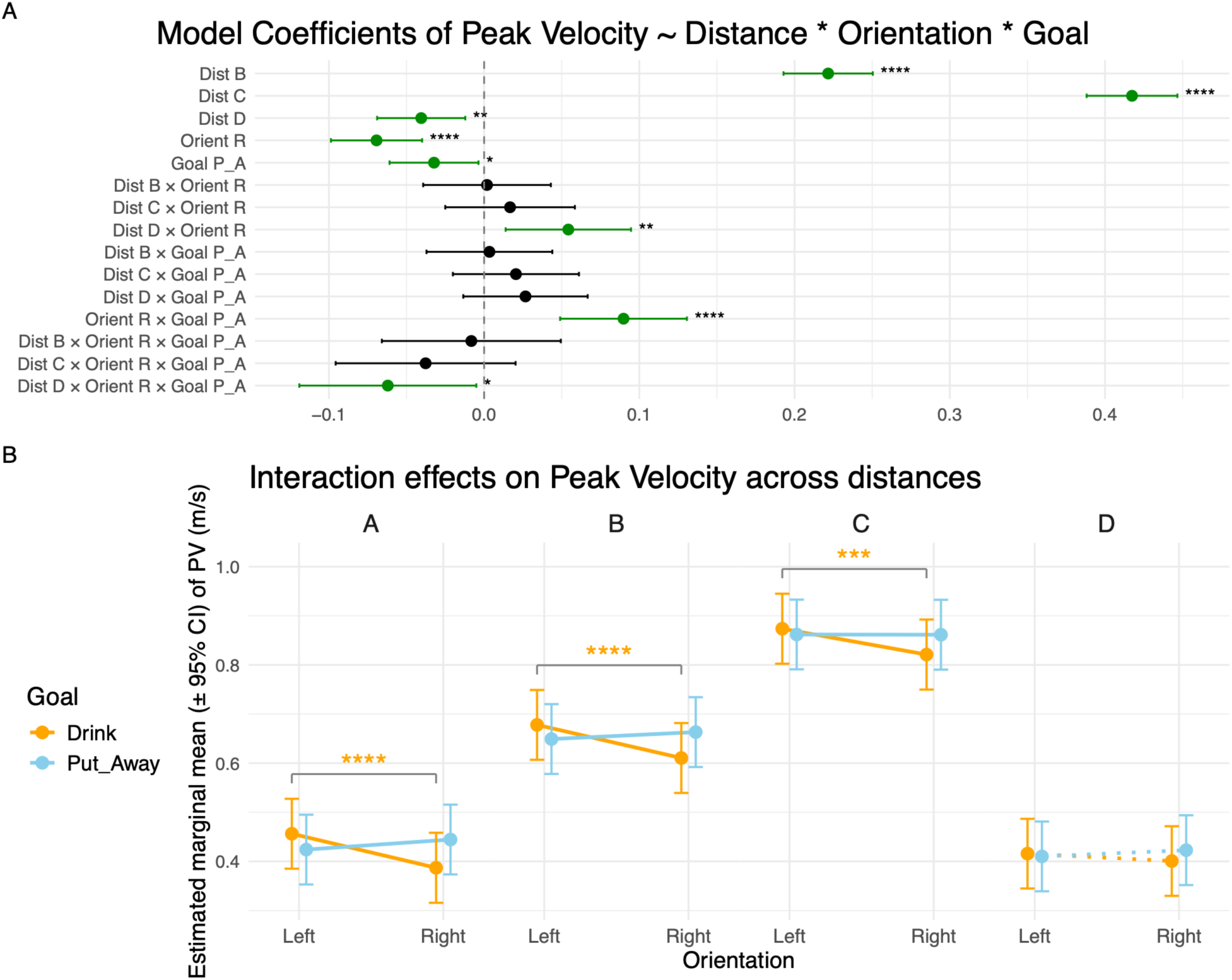
Linear Mixed Model results for Peak Velocity. **A)** Forest plot of the LMM coefficients for Peak Velocity, modelled as a function of Distance, Orientation, and Goal, including all interaction terms. Each row represents one predictor or interaction term, with the point estimate and 95% confidence interval displayed horizontally. Statistically significant coefficients are shown in green; the non-significant coefficient is shown in black. Asterisks denote the level of significance (* p ≤ 0.05, ** p ≤ 0.01, *** p ≤ 0.001, **** p ≤ 0.0001). **B)** Estimated marginal means (± 95% CI) of Peak Velocity as a function of Orientation (Left, Right) at each distance (A, B, C, D), separately for each Goal (Drink: orange; Put Away: sky blue). Brackets with asterisks indicate planned pairwise comparisons between Left and Right orientations within each Distance × Goal combination, with the same significance coding as in panel A. Distance D is shown with dashed lines as it corresponds to a movement directed toward the right corner of the table rather than toward the mug. Note: the **congruent** combinations of Goal and Orientation are represented by Drink-Left and Put Away-Right, the **incongruent** combinations of Goal and Orientation are represented by Drink-Right and Put Away-Left.

**Table 5.** Linear Mixed Model coefficients for Peak Velocity. Fixed-effect estimates from the LMM with Peak Velocity as the dependent variable, modelled as a function of Distance, Orientation, Goal, and all interaction terms. For each predictor, the table reports the coefficient estimate, standard error (SE), degrees of freedom (df), t-value, p-value, and the lower and upper bounds of the 95% confidence interval (CI). Subject ID was included as a random effect. The model intercept corresponds to the combination of Distance = A, Orientation = Left, and Goal = Drink. Distance, Orientation, and Goal are abbreviated as Dist, Orient, and Goal, respectively.

| **Model — PV ~ Distance × Orientation × Goal** |  |  |  |  |  |  |  |
| **Fixed Effect** | **Estimate** | **SE** | **df** | **t** | **p** | **Lower 95% CI** | **Upper 95% CI** |
|---|---|---|---|---|---|---|---|
| (Intercept) | 0.4563 | 0.0363 | 16,5 | 12.572 | **7.24e-10** | 0.3796 | 0.5331 |
| Dist B | 0.2216 | 0.0147 | 28359 | 15.101 | **2.53e-51** | 0.1929 | 0.2504 |
| Dist C | 0.4174 | 0.0149 | 28359 | 27.982 | **5.48e-170** | 0.3882 | 0.4466 |
| Dist D | -0.0405 | 0.0145 | 28360 | -2.798 | **0.0051** | -0.0689 | -0.0121 |
| Orient R | -0.0693 | 0.0150 | 28359 | -4.626 | **3.75e-06** | -0.0987 | -0.0399 |
| Goal P_A | -0.0323 | 0.0146 | 28359 | -2.207 | **0.0273** | -0.0609 | -0.0036 |
| Dist B × Orient R | 0.0019 | 0.0210 | 28359 | 0.089 | 0.9288 | -0.0392 | 0.0430 |
| Dist C × Orient R | 0.0167 | 0.0213 | 28359 | 0.785 | 0.4325 | -0.0250 | 0.0585 |
| Dist D × Orient R | 0.0543 | 0.0206 | 28359 | 2.633 | **0.0085** | 0.0139 | 0.0946 |
| Dist B × Goal P_A | 0.0034 | 0.0207 | 28360 | 0.165 | 0.8688 | -0.0371 | 0.0440 |
| Dist C × Goal P_A | 0.0205 | 0.0207 | 28360 | 0.990 | 0.3224 | -0.0201 | 0.0611 |
| Dist D × Goal P_A | 0.0267 | 0.0204 | 28359 | 1.304 | 0.1924 | -0.0134 | 0.0667 |
| Orient R × Goal P_A | 0.0898 | 0.0208 | 28360 | 4.310 | **1.64e-05** | 0.0490 | 0.1307 |
| Dist B × Orient R × Goal P_A | -0.0082 | 0.0294 | 28360 | -0.280 | 0.7797 | -0.0659 | 0.0494 |
| Dist C × Orient R × Goal P_A | -0.0377 | 0.0296 | 28360 | -1.274 | 0.2026 | -0.0956 | 0.0203 |
| Dist D × Orient R × Goal P_A | -0.0620 | 0.0291 | 28360 | -2.133 | **0.0329** | -0.1190 | -0.0050 |

The main effects of Orientation Right (β = -0.069, *SE* = 0.015, *t*(28359) = -4.626, *p* = 3.75 × 10^-6^, 95% *CI* [-0.099, -0.04]) and Goal Put Away (β = -0.032, *SE* = 0.015, *t*(28359) = -2.207, *p* = 0.027, 95% *CI* [-0.061, -0.004]) were also statistically significant, consistently with movement peak velocity being smaller for movements involving a right-oriented mug compared with a left-oriented mug and for Put Away goal compared with Drink goal.

As well as for the preceding variables (MD, RT), we observed a two-way interaction between Orientation Right and Goal Put Away (that is a congruent combination), resulting in a greater peak velocity (β = 0.09, *SE* = 0.021, *t*(28360) = 4.31, *p* = 1.64 × 10^-5^, 95% *CI* [0.049, 0.131]).

The pairwise comparison (Fig.4B, Table 6) was significant at every reachable distance for the Drink goal, supporting the congruency effect in said condition. The differences between congruency conditions for the goal Drink (Δ = congruent - incongruent) resulted in: **A** (Δ = 0.069, *SE* = 0.015, *z* = 4.626, *p_Bonferroni_* = 3.74 × 10^-6^, 95% *CI* [0.04, 0.099]), **B** (Δ = 0.067, *SE* = 0.015, *z* = 4.594, *p_Bonferroni_* = 4.35 × 10^-6^, 95% *CI* [0.039, 0.096]) and **C** (Δ = 0.053, *SE* = 0.015, *z* = 3.473, *p_Bonferroni_* = 5.0 × 10^-4^, 95% *CI* [0.023, 0.082]).

**Table 6.**
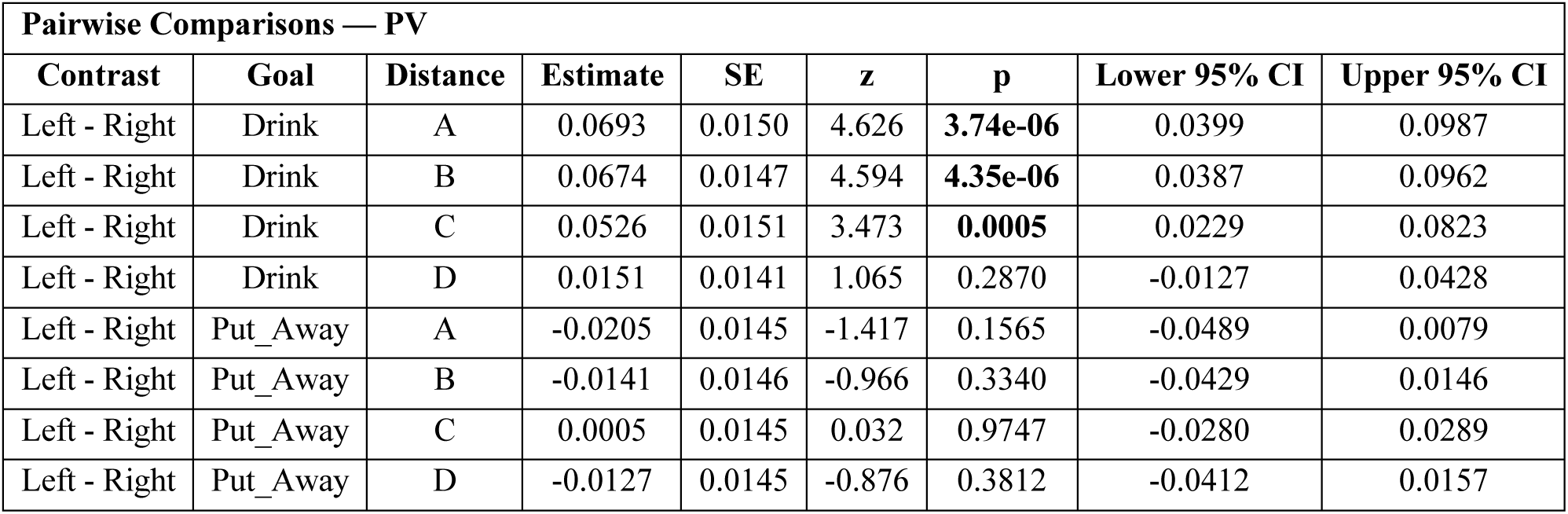
Pairwise comparisons of Peak Velocity between Left and Right orientations. Estimated marginal mean differences (Δ = Left − Right) between orientation conditions at each Distance × Goal combination (2 Goals × 4 Distances), derived from the LMM using the emmeans package. P-values are Bonferroni-corrected for multiple comparisons. Note: For the Drink goal, Left − Right corresponds to Δ = congruent − incongruent; for the Put Away goal, Left − Right corresponds to Δ = incongruent − congruent.

| **Pairwise Comparisons — PV** |  |  |  |  |  |  |  |  |
| **Contrast** | **Goal** | **Distance** | **Estimate** | **SE** | **z** | **p** | **Lower 95% CI** | **Upper 95% CI** |
|---|---|---|---|---|---|---|---|---|
| Left - Right | Drink | A | 0.0693 | 0.0150 | 4.626 | **3.74e-06** | 0.0399 | 0.0987 |
| Left - Right | Drink | B | 0.0674 | 0.0147 | 4.594 | **4.35e-06** | 0.0387 | 0.0962 |
| Left - Right | Drink | C | 0.0526 | 0.0151 | 3.473 | **0.0005** | 0.0229 | 0.0823 |
| Left - Right | Drink | D | 0.0151 | 0.0141 | 1.065 | 0.2870 | -0.0127 | 0.0428 |
| Left - Right | Put_Away | A | -0.0205 | 0.0145 | -1.417 | 0.1565 | -0.0489 | 0.0079 |
| Left - Right | Put_Away | B | -0.0141 | 0.0146 | -0.966 | 0.3340 | -0.0429 | 0.0146 |
| Left - Right | Put_Away | C | 0.0005 | 0.0145 | 0.032 | 0.9747 | -0.0280 | 0.0289 |
| Left - Right | Put_Away | D | -0.0127 | 0.0145 | -0.876 | 0.3812 | -0.0412 | 0.0157 |

In summary, we observed a congruency effect within the reachable space for the Drink goal but no effect outside the participants’ reach, supporting the conclusion that the congruency effect is conditional on the perceived possibility of interacting with the object.

### Log-dimensionless-jerk

To operationalize congruency beyond a qualitative definition, we tested whether movement smoothness, assessed through a Log-dimensionless-jerk (LDJ) index, was sensitive to the congruency manipulation, expecting jerkier and hence more costly movements (Wong et al., 2021) in the incongruent condition. In brief, this metric may be interpreted as a proxy of metabolic cost if we assume that this cost is a force-rate function during reaching. In other words, F’=m×jerk. Given that our design gathered kinematic measurements only, a direct recording/computation of force-rate (i.e., the time derivative of muscle force) was not feasible. Therefore, our analysis was limited to jerk, quantified using the LDJ metric proposed by (Melendez-Calderon et al., 2021). In the present work we changed the sign of the LDJ metric as compared with the original work only for ease of interpretation, where LDJ equals zero for completely smooth movements and increases as the trajectory becomes jerkier. Related negative model coefficients would indicate that the variable is associated with an increase in movement smoothness and, conversely otherwise.

The LMM on LDJ (Fig 5A, Table 7) revealed a statistically significant main effect of Distance, whereby smoothness increased with distance: **B** (β = -0.307, *SE* = 0.079, *t*(26956) = -3.867, *p* = 1.1× 10^-4^, 95% *CI* [-0.463, -0.152]), **C** (β = -0.683, *SE* = 0.081, *t*(26956) = -8.422, *p* = 3.88 × 10^-17^, 95% *CI* [-0.842, -0.524]) and **D** (β = -1.829, *SE* = 0.078, *t*(26956) = -23.32, *p* = 4.2 × 10^-119^, 95% *CI* [-0.842, -0.524]). Distance D required a different movement, directed toward the right corner of the table, and therefore exhibited greater smoothness than all other conditions, as expected, and did not differ between goals, as the same movement was executed regardless of goal, as illustrated in Fig 5B.

**Figure 5.**
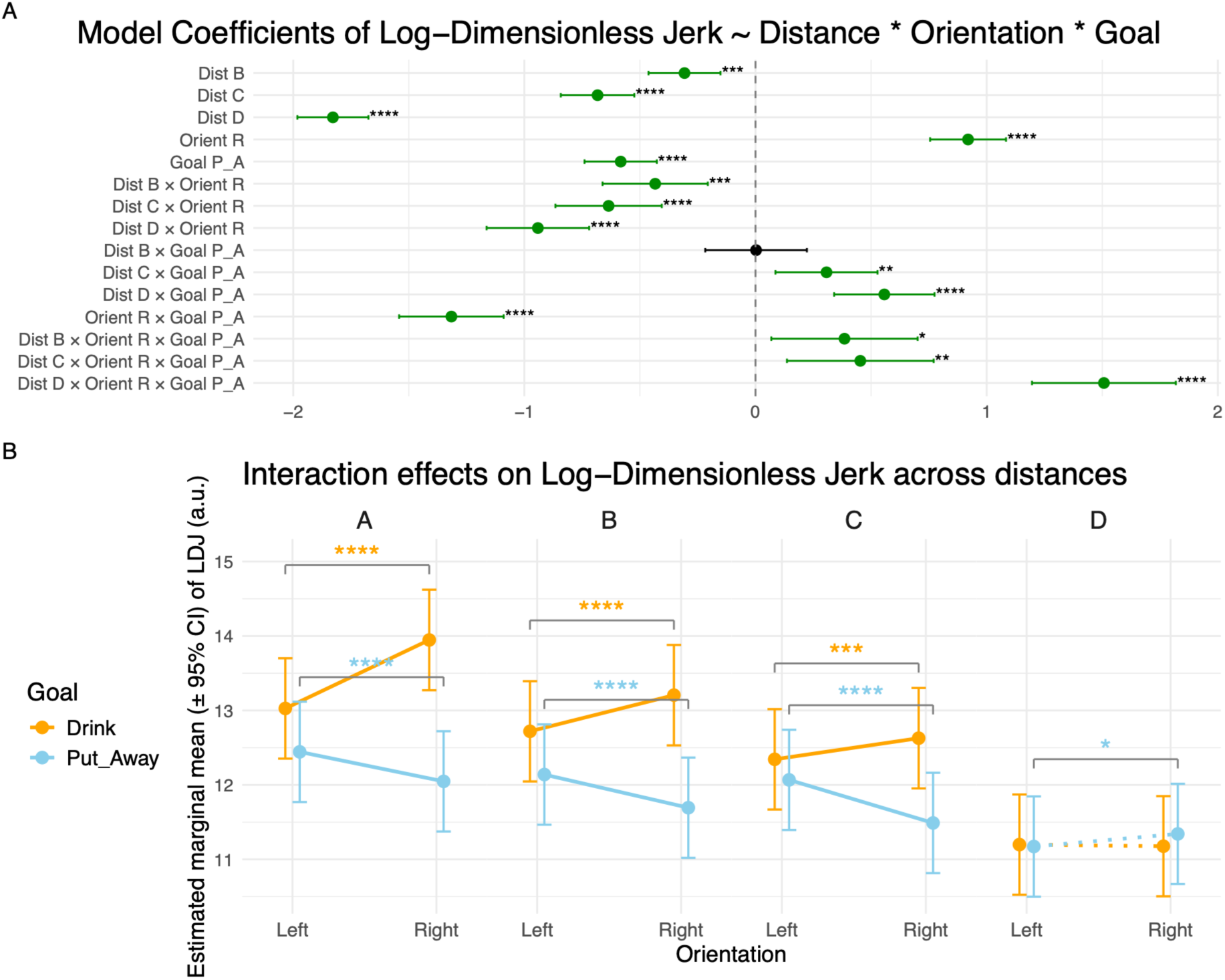
Linear Mixed Model results for Log-Dimensionless Jerk. **A.** Forest plot of the LMM coefficients for Log-Dimensionless Jerk, modelled as a function of Distance, Orientation, and Goal, including all interaction terms. Each row represents one predictor or interaction term, with the point estimate and 95% confidence interval displayed horizontally. Statistically significant coefficients are shown in green; the non-significant coefficient is shown in black. Asterisks denote the level of significance (* p ≤ 0.05, ** p ≤ 0.01, *** p ≤ 0.001, **** p ≤ 0.0001). **B.** Estimated marginal means (± 95% CI) of Log-Dimensionless Jerk as a function of Orientation (Left, Right) at each distance (A, B, C, D), separately for each Goal (Drink: orange; Put Away: sky blue). Brackets with asterisks indicate planned pairwise comparisons between Left and Right orientations within each Distance × Goal combination, with the same significance coding as in panel A. Distance D is shown with dashed lines as it corresponds to a movement directed toward the right corner of the table rather than toward the mug. Note: the **congruent** combinations of Goal and Orientation are represented by Drink-Left and Put Away-Right, the **incongruent** combinations of Goal and Orientation are represented by Drink-Right and Put Away-Left.

**Table 7.** Linear Mixed Model coefficients for Log-Dimensionless Jerk. Fixed-effect estimates from the LMM with Log-Dimensionless Jerk as the dependent variable. modelled as a function of Distance, Orientation, Goal, and all interaction terms. For each predictor, the table reports the coefficient estimate, standard error (SE). degrees of freedom (df), t-value, p-value, and the lower and upper bounds of the 95% confidence interval (CI). Subject ID was included as a random effect. The model intercept corresponds to the combination of Distance = A, Orientation = Left, and Goal = Drink, Distance, Orientation, and Goal are abbreviated as Dist. Orient, and Goal, respectively.

| **Model — LDJ ~ Distance × Orientation × Goal** |  |  |  |  |  |  |  |
|---|---|---|---|---|---|---|---|
| **Fixed Effect** | **Estimate** | **SE** | **df** | **t** | **p** | **Lower 95% CI** | **Upper 95% CI** |
| (Intercept) | 13.0272 | 0.3438 | 14,7 | 37.888 | **4.40e-16** | 12.2932 | 13.7613 |
| Dist B | -0.3071 | 0.0794 | 26956 | -3.867 | **0.0001** | -0.4628 | -0.1515 |
| Dist C | -0.6833 | 0.0811 | 26956 | -8.422 | **3.88e-17** | -0.8424 | -0.5243 |
| Dist D | -1.8285 | 0.0784 | 26956 | -23.320 | **4.20e-119** | -1.9822 | -1.6748 |
| Orient R | 0.9195 | 0.0838 | 26956 | 10.974 | **5.85e-28** | 0.7553 | 1.0838 |
| Goal P_A | -0.5832 | 0.0798 | 26956 | -7.307 | **2.81e-13** | -0.7397 | -0.4268 |
| Dist B × Orient R | -0.4340 | 0.1162 | 26956 | -3.734 | **0.0002** | -0.6618 | -0.2062 |
| Dist C × Orient R | -0.6356 | 0.1174 | 26956 | -5.415 | **6.18e-08** | -0.8656 | -0.4055 |
| Dist D × Orient R | -0.9415 | 0.1132 | 26956 | -8.317 | **9.41e-17** | -1.1634 | -0.7196 |
| Dist B × Goal P_A | 0.0025 | 0.1122 | 26956 | 0.022 | 0.9825 | -0.2174 | 0.2223 |
| Dist C × Goal P_A | 0.3071 | 0.1127 | 26956 | 2.726 | **0.0064** | 0.0863 | 0.5279 |
| Dist D × Goal P_A | 0.5572 | 0.1108 | 26956 | 5.028 | **4.98e-07** | 0.3400 | 0.7745 |
| Orient R × Goal P_A | -1.3158 | 0.1154 | 26956 | -11.405 | **4.61e-30** | -1.5419 | -1.0897 |
| Dist B × Orient R × Goal P_A | 0.3849 | 0.1615 | 26956 | 2.383 | **0.0172** | 0.0683 | 0.7015 |
| Dist C × Orient R × Goal P_A | 0.4534 | 0.1620 | 26956 | 2.799 | **0.0051** | 0.1359 | 0.7708 |
| Dist D × Orient R × Goal P_A | 1.5072 | 0.1587 | 26956 | 9.496 | **2.35e-21** | 1.1961 | 1.8182 |

The main effects of Orientation (β = 0.92, *SE* = 0.084, *t*(26956) = 10.974, *p* = 5.85 × 10^-28^, 95% *CI* [0.755, 1.084]) and Goal (β = -0.583, *SE* = 0.08, *t*(26956) = -7.307, *p* = 2.81 × 10^-13^, 95% *CI* [-0.74, -0.427]) were also statistically significant, consistently with movements involving a right-oriented mug were less smooth than those directed toward a left-oriented mug. Movement smoothness also differed between goals, being greater for the Put Away goal than for the Drink goal.

The interaction term was statistically significant, accordingly with the aforementioned variables (MD, RT, PV), between Orientation Right and Goal Put Away (β = -1.316, *SE* = 0.115, *t*(26956) = -11.405, *p* = 4.61 × 10^-30^, 95% *CI* [-1.542, -1.09]), indicating smoother movement for congruent conditions.

The pairwise comparison (Fig 5B, Table 8) revealed that all differences between congruent and incongruent conditions were significant, with the exception of distance D in Drink trials. For the Drink goal, the differences (Drink Δ = congruent - incongruent) for the four distances were: **A** (Δ = - 0.92, *SE* = 0.084, *z* = -10.974, *p_Bonferroni_* = 5.1 × 10^-28^, 95% *CI* [-1.084, -0.755]), **B** (Δ = -0.487, *SE* = 0.081, *z* = -6.031, *p_Bonferroni_* = 1.627 × 10^-9^, 95% *CI* [-0.643, -0.328]), **C** (Δ = -0.284, *SE* = 0.082, *z* = -3.455, *p_Bonferroni_* = 5.0 × 10^-4^, 95% *CI* [-0.445, -0.123]) and **D** (Δ = 0.022, *SE* = 0.076, *z* = 0.289, *p_Bonferroni_* = 0.773, 95% *CI* [-0.127, 0.171]). For the Put Away goal, the differences (Put Away Δ = incongruent - congruent) for the four distances were: **A** (Δ = 0.396, *SE* = 0.079, *z* = 5.001, *p_Bonferroni_* = 5.694 × 10^-28^, 95% *CI* [0.241, 0.552]), **B** (Δ = 0.445, *SE* = 0.079, *z* = 5.624, *p_Bonferroni_* = 1.861× 10^-8^, 95% *CI* [0.29, 0.601]), **C** (Δ = 0.579, *SE* = 0.078, *z* = 7.38, *p_Bonferroni_* = 1.580 × 10^-13^, 95% *CI* [0.425, 0.732]) and **D** (Δ = -0.169, *SE* = 0.078, *z* = -2.175, *p_Bonferroni_* = 0.029, 95% *CI* [-0.322, 0.017]).

**Table 8.** Pairwise comparisons of Log-Dimensionless Jerk between Left and Right orientations. Estimated marginal mean differences (Δ = Left − Right) between orientation conditions at each Distance × Goal combination (2 Goals × 4 Distances), derived from the LMM using the *emmeans* package. P-values are Bonferroni-corrected for multiple comparisons. Note: For the Drink goal, Left − Right corresponds to Δ = congruent − incongruent; for the Put Away goal, Left − Right corresponds to Δ = incongruent − congruent.

| **Pairwise Comparisons — LDJ** |  |  |  |  |  |  |  |  |
|---|---|---|---|---|---|---|---|---|
| **Contrast** | **Goal** | **Distance** | **Estimate** | **SE** | **z** | **p** | **Lower 95% CI** | **Upper 95% CI** |
| Left - Right | Drink | A | -0.9195 | 0.0838 | -10.974 | **5.10e-28** | -1.0838 | -0.7553 |
| Left - Right | Drink | B | -0.4856 | 0.0805 | -6.031 | **1.63e-09** | -0.6433 | -0.3278 |
| Left - Right | Drink | C | -0.2840 | 0.0822 | -3.455 | **0.0005** | -0.4450 | -0.1229 |
| Left - Right | Drink | D | 0.0220 | 0.0761 | 0.289 | 0.7727 | -0.1272 | 0.1712 |
| Left - Right | Put_Away | A | 0.3963 | 0.0792 | 5.001 | **5.69e-07** | 0.2410 | 0.5516 |
| Left - Right | Put_Away | B | 0.4454 | 0.0792 | 5.624 | **1.86e-08** | 0.2902 | 0.6005 |
| Left - Right | Put_Away | C | 0.5785 | 0.0784 | 7.380 | **1.58e-13** | 0.4248 | 0.7321 |
| Left - Right | Put_Away | D | -0.1694 | 0.0779 | -2.175 | **0.0297** | -0.3221 | -0.0167 |

In summary, the qualitative definition of congruency is supported by LDJ quantitative measure of physical effort and motor cost. Congruent movements were smoother (i.e., required less effort) than incongruent ones.

### Two alternatives forced choice task

We finally assessed whether participants were sensitive to physical effort costs in our setup – by asking them to make a series of binary decisions between higher or lower effort options (goal-orientation combinations) presented on a computer screen. The analysis of preferences has been conducted at first by comparing the proportion of times one option (the one on the left, counterbalancing their position across trials) was chosen over the total number of times the option was displayed to the participant as shown in Table 9. To test that the proportion of times the option on the left has been chosen more than 50% of times, a two-tailed binomial test has been conducted, subsequently correcting the p-values for multiple comparisons with the Benjamini-Hochberg correction (Benjamini & Hochberg, 1995).

**Table 9.** Pairwise preference proportions from the 2AFC task. For each pair of options presented, the table reports the proportion of trials in which the first-listed option was chosen over the second-listed one, computed as #first / (#first + #second). Statistical significance was assessed using a one-sample proportion test against chance level (0.5), with p-values corrected for multiple comparisons using the Benjamini-Hochberg procedure.

| **Offer** | **Proportion of chosen first option** | **Benjamini-Hochberg corrected p-values** |
|---|---|---|
| Drink Left vs Drink Right | 0.94 | 2.70 × 10<sup>-62</sup> |
| Drink Left vs Put Away Left | 0.83 | 3.72 × 10<sup>-31</sup> |
| Put Away Right vs Drink Right | 0.94 | 1.69 × 10<sup>-60</sup> |
| Put Away Right vs Put Away Left | 0.85 | 1.58 × 10<sup>-36</sup> |
| Drink Left vs Put Away Right | 0.63 | 6.79 × 10<sup>-6</sup> |
| Drink Right vs Put Away Left | 0.29 | 1.61 × 10<sup>-12</sup> |

Subsequently, we implemented a logistic regression (Equation 3) fitted through the *lme4 (Bates et al., 2015)* R package to predict the probability of choosing one option (the left one) given the log ratio of the LDJ values of the two options (the left option divided by the right option), computed by subject as the average LDJ for the mug state and action combination recorded during the action experiment. The main effect of the logistic regression was significant (β_1_ = -8.437, *SE* = 0.838, *z* = −10.063, *p* = 2.0 × 10^-16^, 95% *CI* [-10.102, -6.814]), meaning that the group preferred to choose the smoother (or less costly) movement option. The intercept was not significant (β_0_ = −0.032, *SE* = 0.049, *z* = −0.654, *p* = 0.513, 95% *CI* [-0.128, 0.064]) meaning that no bias toward the left side option was present (Fig 6, Table 10).

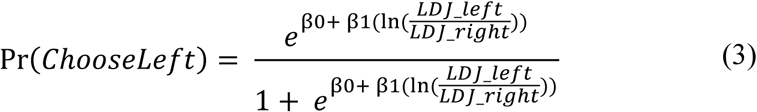

**Figure 6.**
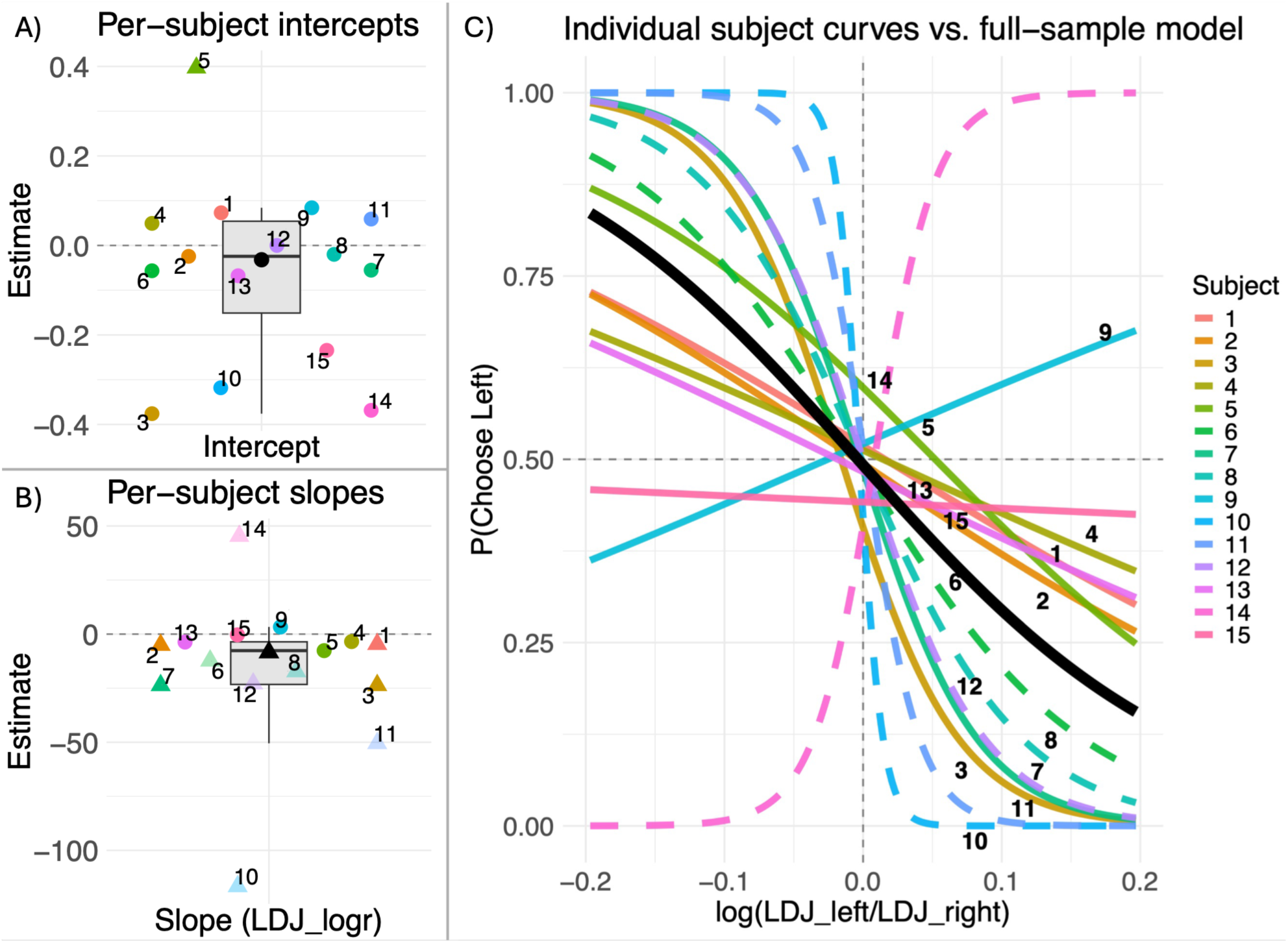
Two alternatives forced choice task modelling. **A.** Subjects’ intercept estimates in coloured symbols accordingly with the legend. The triangle indicates a p-value for the coefficient <.05, the circle otherwise. The black symbol represents the full logistic regression estimate. **B.** Subjects’ slope estimates in coloured symbols accordingly with the legend. The triangle indicates a p-value for the coefficient <.05, the circle otherwise. The black symbol represents the full logistic regression estimate. **C.** Individual logit functions coloured accordingly with the legend shown alongside the full logistic regression fit. **Note**: Faded dots/triangles and dashed lines represent subjects for which the individual logistic regression resulted in a SE > 5 for either the intercept or the slope coefficient, denoting high uncertainty of the estimate.

**Table 10.**
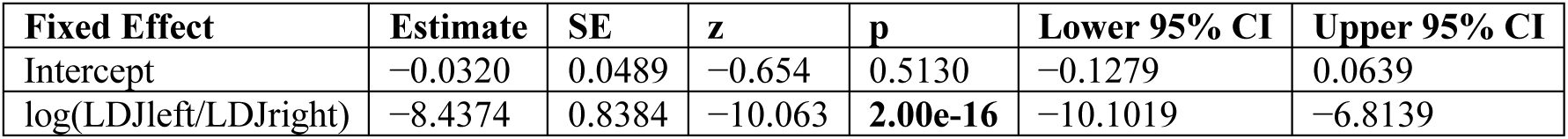
Logistic Regression (GLM) to model preference given log-dimensionless jerk left and right values ratio related to the options.

## Discussion

Seeing an object elicits motor processes and behaviours congruent with the object affordances (Tucker & Ellis, 1998, 2001). In this work, we investigated (1) whether the congruency effect (i.e., the facilitation of the motor response elicited by the compatibility between the effector used to report the response and the stimulus) was present in naturalistic, goal-directed actions with a physical object (a mug) during simulated functional use (drinking from it or putting it away); (2) whether this effect was limited to the reachable space; (3) whether a physical-effort metric derived from arm kinematics (log-dimensionless jerk, LDJ) could account for the congruency effect; and (4) whether participants preferred movements associated with lower physical effort, as quantified by this metric.

We report four main findings. First, the results of the main Linear Mixed Model (LMM), through the Orientation Right × Goal Put Away interaction term, and the planned pairwise comparisons between congruency conditions at each distance support the conclusion that the congruency effect emerges across different indicators. Congruency influenced Movement Duration (MD): congruent movements were shorter than incongruent ones, an effect that might reflect lower movement complexity. Furthermore, congruency effects emerged for both the Put Away and the Drink goals, although in different variables: Reaction Time (RT) and Peak Velocity (PV), respectively (see also the Supplementary Materials for a congruency effect on absolute time to peak velocity for the Drink goal). RT was shorter in the congruent condition, but only for the Put Away goal and at reachable distances. It is plausible that the longer RT for incongruent stimuli reflected the longer planning time required to review or adjust the affordance evoked for a canonical interaction and the longer movement duration (as supported by the MD result above), and the longer trajectory involved in the movement. However, the latter interpretation is not supported by our results on trajectory length (see Supplementary Material), which showed no difference between congruent and incongruent trajectories in the Put Away goal, the goal for which we observed a congruency effect on RT. Another possible explanation is that eliciting the congruency effect requires the goal–action–object coupling to be unusual rather than automatic. Our data show that the Drink goal was highly learned and automatic (as reflected in faster RTs), which may explain the absence of a congruency effect on RT for this goal. Conversely, PV displayed a congruency effect for the Drink goal at reachable distances, with higher PV in the congruent than in the incongruent condition. Interestingly, although the effects on RT and PV were not observed simultaneously for the same goal, taken separately they parallel the findings on movement vigour (Summerside et al., 2018), which showed that movements toward better-rewarded goals were executed with shorter RTs and higher PVs. In our experiment, reward was not manipulated, but the physical effort of executing an action was hypothesized to be higher in the incongruent than in the congruent condition (as supported by our LDJ characterization of movement and discussed further below). This higher effort may have led the more costly conditions to be perceived as less desirable (as observed in the two-alternative forced-choice task). Hence, behaviour may have been affected in both experiments by economic assessments of movement. Why the congruency effect was observed in different variables for different goals remains an open question.

Second, importantly, the congruency effects on both RT and PV were limited to the reachable space, which in this experiment coincides with the space that allows object interaction. This is consistent with peri-personal space studies addressing the effect of action feasibility on affordance evocation (Costantini et al., 2010). In addition, this result supports the idea that a covert affordance/motor simulation mechanism was engaged during the action task and, therefore, that the observed patterns are not merely related to the visual properties of the stimulus.

Third, the congruency effect was present for both goals in the LDJ analysis: congruent movements had smoother trajectories than incongruent ones within each goal. This metric of movement smoothness can be interpreted as a proxy for physical effort and/or as a quantitative description of movement complexity. Its pattern was consistent with the qualitative expectation that congruent movements are less demanding.

Finally, our results show that, when given a choice between two options (2AFC task), participants preferred congruent over incongruent combinations. A logistic regression using LDJ values (as a proxy for physical effort) to predict the probability of choosing an option showed that preference leaned toward the least effortful option, with no bias toward the left or right position of the option. Given that the reward for participating in the experiment was the same for all participants, this last result is consistent with studies on effort-based decision-making (Hartmann et al., 2013; Matthews et al., 2023; Morel et al., 2017; Skvortsova et al., 2014), which frame motor cost as a discounting factor that decreases the net utility of an option.

Together, these results also show that distance D (which required moving the right hand to the right corner of the table) was associated with both the shortest MD and the greatest smoothness, possibly reflecting a lower demand in the execution of the associated movement. Surprisingly, however, PV was the lowest and RT the longest among all distances. It is possible that participants expected to execute a forward movement toward the mug, as opposed to a sideways movement to the right corner of the table, at the unreachable distance D and in catch trials without the mug (i.e., the No-Go condition). If this expectation is built through experience, the probability of a forward movement should be higher than that of its complementary event, a sideways movement to the right. Indeed, the cumulative probabilities of the three forward-movement conditions (the reachable distances A, B, and C) and of the two "move sideways to the right" conditions (distance D and the No-Go catch trials) were three out of five and two out of five, respectively. Hence, it is possible that the expected low-level movement pattern was to move the arm forward, as this was the most likely event, and that, as a consequence, the longer RTs and lower PVs at distance D reflected the cost of changing the movement rule (i.e., moving right instead of forward).

This effect might operate at a smaller scale, thereby explaining, at least in part, the congruency effect as the amount of divergence from a prior expectation (smaller for congruent than for incongruent stimuli). This possibility could be tested with a paradigm that compares congruent and incongruent stimuli (in a motor context) while keeping constant the physical effort and the familiarity with the goal-directed actions across congruency conditions, and while manipulating the likelihood of each goal-directed action, in order to assess to what extent the congruency effect entails a process of adjustment of movement beliefs.

An unexpected result was the inversion of the RT pattern at distance B for the Put Away goal, where the congruent condition was slower than the incongruent one. Notably, distance B was the fastest and qualitatively the most ergonomic among the reachable distances: distance A was very close to the chin rest, whereas distance C was at the boundary of the reachable space. This result is difficult to reconcile with the idea that the congruency effect necessarily reflects automatic processes, because the only distance at which a reversed pattern was observed was the one with the shortest RTs. Moreover, the LDJ analysis showed the predicted pattern of smoother trajectories for congruent stimuli, which makes this result even more difficult to interpret. The present experiment does not allow us to explain this pattern, and further investigation is required.

Another unexpected result was the statistically significant difference between congruency conditions in the Put Away goal at the unreachable distance D, for the MD and LDJ variables: the congruent condition was both longer and less smooth, therefore showing a reverse pattern relative to the other distances. It is possible that inhibiting the reach movement (owing to the impossibility of reaching the object and the instruction to move sideways to the right in that condition) was more demanding given the possibly stronger mental representation of the congruent combination.

Altogether, from a methodological perspective, these results also highlight the importance of including kinematic variables when investigating this process, since the congruency effect has been consistently observed through different markers: MD, RT (in the Put Away goal), PV (in the Drink goal), and LDJ.

The present study contributes to a large body of research investigating how object affordances are processed. This literature has shown that objects can automatically evoke action possibilities, even when the actions they afford are irrelevant to the task at hand (Ellis & Tucker, 2000; Tucker & Ellis, 1998, 2001). This phenomenon is typically reflected in faster reaction times when stimulus features are congruent with the required response, an effect commonly referred to as the congruency effect. Importantly, such effects appear to be constrained by the agent’s perceived action capabilities, emerging only within the space in which an interaction with the object is considered achievable (Costantini et al., 2010, 2011; Kalénine et al., 2016). More broadly, it has been proposed that action selection in the brain involves a competition between multiple simultaneously represented affordances, i.e., the “affordance competition” hypothesis (Cisek, 2007; Cisek & Kalaska, 2010; Pezzulo & Cisek, 2016) and that this dynamical competition produces a rich interplay between decisions and actions (Araújo et al., 2006; Barca & Pezzulo, 2012; Cisek & Pastor-Bernier, 2014; Cos et al., 2021; Djebbara et al., 2019; Gordon et al., 2021; Grießbach et al., 2021; Marcos et al., 2015; Pastor-Bernier & Cisek, 2011; Thura & Cisek, 2014; Wispinski et al., 2018). The main contribution of this study is showing that object–action congruency shapes the execution of both naturalistic goal-directed movements and action preferences and that these effects are specific to reachable space. Furthermore, our study provides a quantitative measure of congruency – movement smoothness – that helps understanding deep connections between congruency effects and efficient motor execution, therefore linking to a long tradition that highlights efficiency as a key principle for decision-making and motor control (Gallivan et al., 2018; Lepora & Pezzulo, 2015; Parr et al., 2022; Priorelli et al., 2025; Shadmehr & Mussa-Ivaldi, 2012; Trommershäuser et al., 2008).

Although this design has the advantage of using real-object actions to simulate a more ecological interaction than previous experiments, the study also has some limitations. First, the horizontal position of the mug might have been perceived as unusual and unrelated to its typical function, potentially preventing participants from experiencing a truly congruent stimulus, that is, one that is both commonly experienced (e.g., an upward-oriented mug) and aligned with the effector. However, several studies (Borghi & Riggio, 2009; Bub et al., 2015) suggest that viewing an object elicits features from multiple modalities, including its canonical state and function, regardless of its depicted state. If this is the case, the congruency effect on RT observed here might result from the difference between the action that would be chosen for the object in its canonical state and the action required for the object in its current state. This interpretation would extend the discrete concept of congruency to a continuous space of adaptation from expected to observed affordances. Including a canonical condition would be necessary to test this hypothesis. A surprising result in our experiment is consistent with this interpretation: despite the greater movement smoothness for the Put Away goal, RTs were longer than for the Drink goal. To reconcile this result with the proposed interpretation, we suggest that the additional time needed to initiate movement in the Put Away condition might reflect a conflict between the cued goal and the canonical function elicited by the object. Including unlikely goal–object combinations in future studies could help test this prediction.

A second limitation concerns the possibility of interpreting the presence of the congruency effect as a function of interaction feasibility. Feasibility was manipulated in our task by including an unreachable distance (D). Because the object was unreachable, a response different from reaching it was required, which in our case consisted of touching the right corner of the table. Hence, we cannot exclude that the absence of a congruency effect at this unreachable distance is due to the process of changing the response rule (i.e., moving to the right instead of forward toward the mug). This intrinsic limitation of a design requiring overt, naturalistic goal-directed action might be addressed through a control experiment exploiting a response modality that is independent of the object’s reachability, similar to studies that investigated the congruency effect through pantomiming an action toward a screen (Costantini et al., 2010).

## Acknowledgements

This research received funding from the European Research Council under the Grant Agreement No. 820213 (ThinkAhead) and the Ministry of University and Research, PRIN 20229Z7M8N for RM and GP; and from the Spanish Ministry of Science, Innovation and Universities Grant with PID2023-149210NB-I00 for IC and MH. We used a Generative AI model to correct typographical errors and edit language for clarity.

## Supplementary Material

### Handedness

In Table 1S we report the results of the Edinburgh Handedness Inventory as the Laterality Quotient (LQ) for teach participant (ID). The majority of participants have a positive score, indicating a tendency toward right handedness, with a mean of 59.25 ± 28.84 std.

**Table 1S.**

| ID | 1 | 2 | 3 | 4 | 5 | 6 | 7 | 8 | 9 | 10 | 11 | 12 | 13 | 14 | 15 |
|---|---|---|---|---|---|---|---|---|---|---|---|---|---|---|---|
| LQ | 61.90 | 80.95 | 66.67 | 71.43 | 100.00 | 26.32 | -6.67 | 87.50 | 68.42 | 25.00 | 55.56 | 73.91 | 50.00 | 90.91 | 36.84 |

### Movement trajectory length

**Figure 1S.**
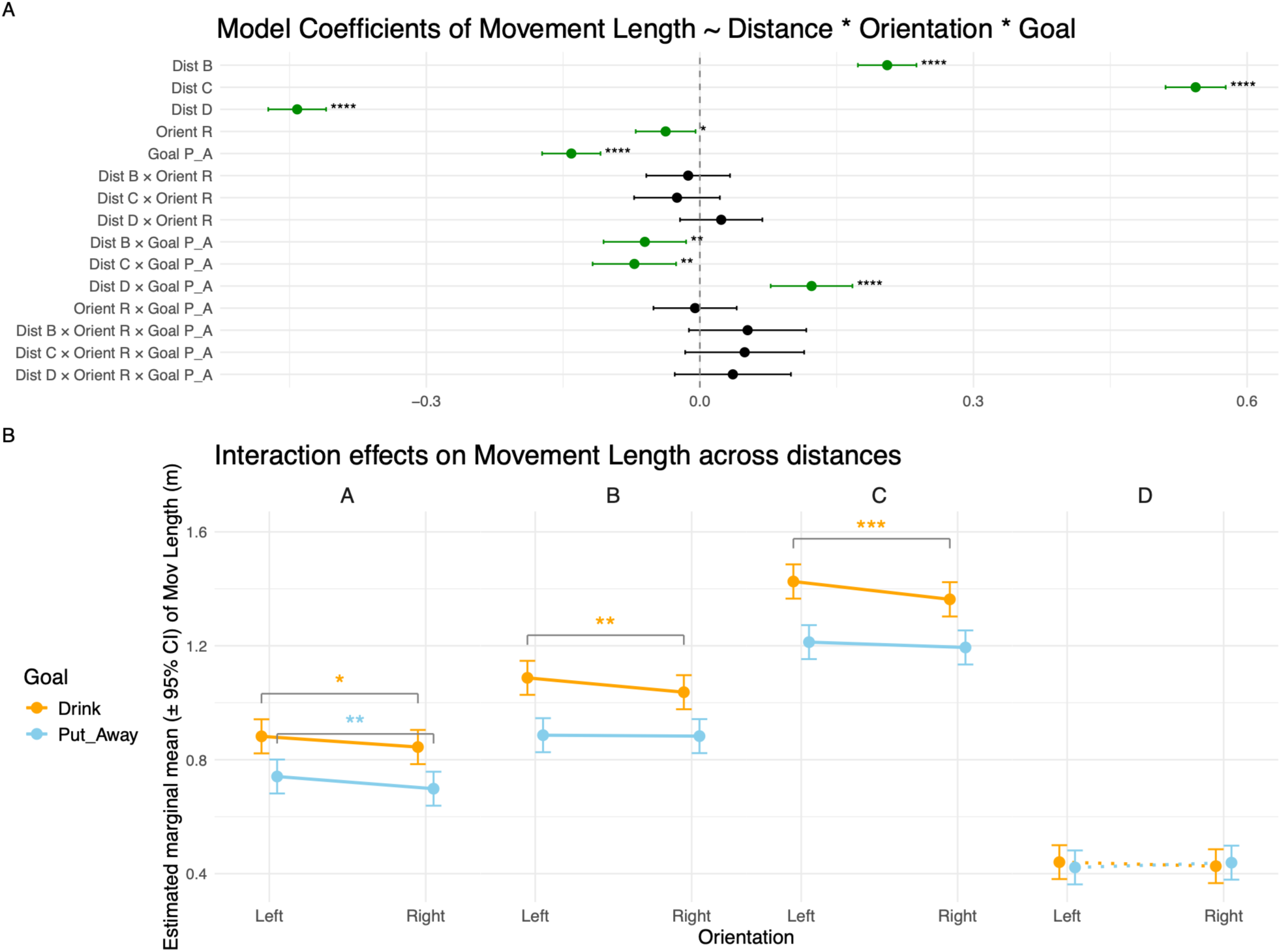
Linear Mixed Model results for Movement Length. **A)** Forest plot of the LMM coefficients for Movement Length, modelled as a function of Distance, Orientation, and Goal, including all interaction terms. Each row represents one predictor or interaction term, with the point estimate and 95% confidence interval displayed horizontally. Statistically significant coefficients are shown in green; the non-significant coefficient is shown in black. Asterisks denote the level of significance (* p ≤ 0.05, ** p ≤ 0.01, *** p ≤ 0.001, **** p ≤ 0.0001). **B)** Estimated marginal means (± 95% CI) of Movement Length as a function of Orientation (Left, Right) at each distance (A, B, C, D), separately for each Goal (Drink: orange; Put Away: sky blue). Brackets with asterisks indicate planned pairwise comparisons between Left and Right orientations within each Distance × Goal combination, with the same significance coding as in panel A. Distance D is shown with dashed lines as it corresponds to a movement directed toward the right corner of the table rather than toward the mug. Note: the **congruent** combinations of Goal and Orientation are represented by Drink-Left and Put Away-Right, the **incongruent** combinations of Goal and Orientation are represented by Drink-Right and Put Away-Left.

**Table 2S.**
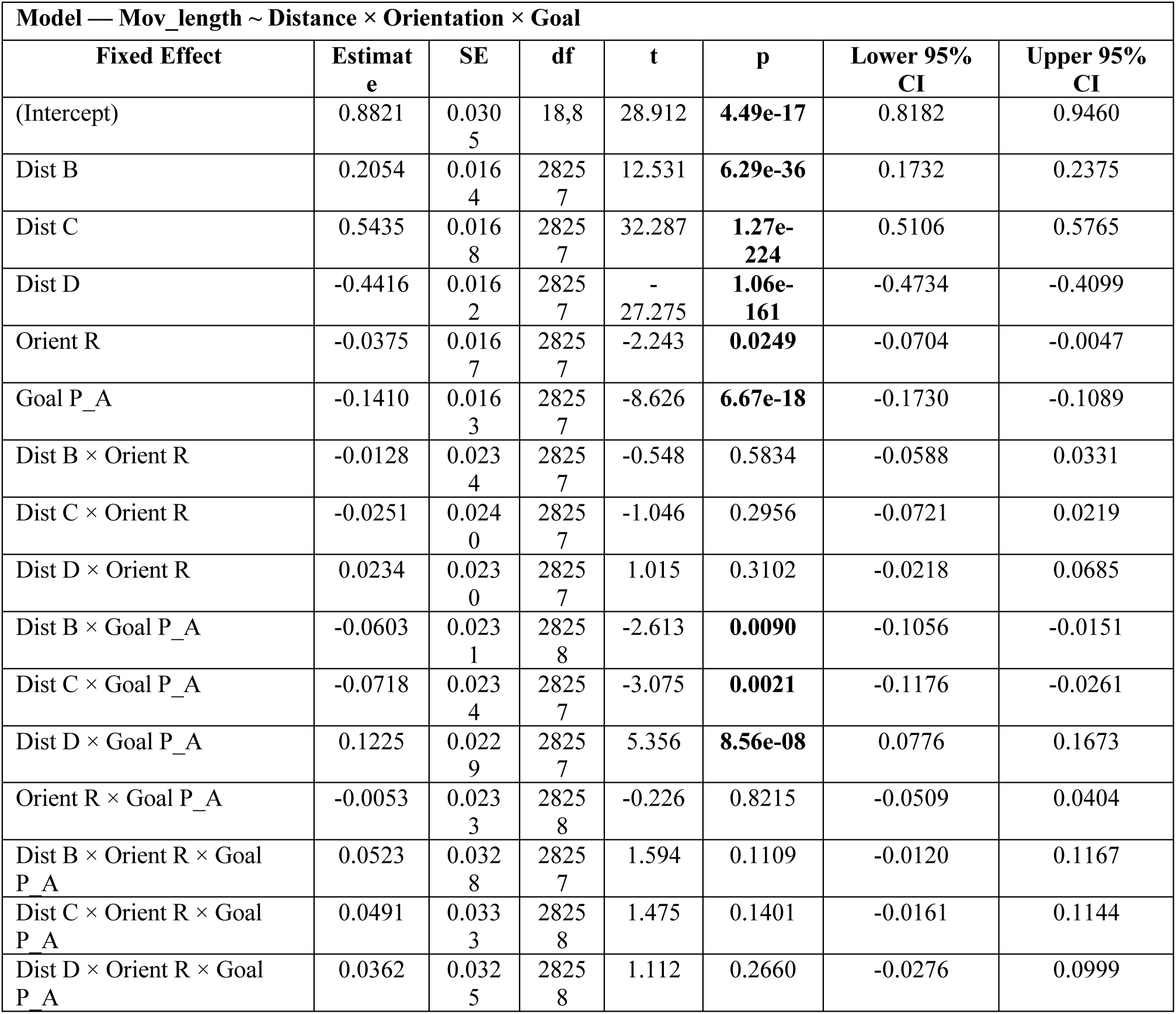
Linear Mixed Model coefficients for Movement Length. Fixed-effect estimates from the LMM with Movement Length as the dependent variable, modelled as a function of Distance, Orientation, Goal, and all interaction terms. For each predictor, the table reports the coefficient estimate, standard error (SE), degrees of freedom (df), t-value, p-value, and the lower and upper bounds of the 95% confidence interval (CI). Subject ID was included as a random effect. The model intercept corresponds to the combination of Distance = A, Orientation = Left, and Goal = Drink. Distance, Orientation, and Goal are abbreviated as Dist, Orient, and Goal, respectively.

**Table 3S.** Pairwise comparisons of Movement Length between Left and Right orientations. Estimated marginal mean differences (Δ = Left − Right) between orientation conditions at each Distance × Goal combination (2 Goals × 4 Distances), derived from the LMM using the emmeans package. P-values are Bonferroni-corrected for multiple comparisons. Note: For the Drink goal, Left − Right corresponds to Δ = congruent − incongruent; for the Put Away goal, Left − Right corresponds to Δ = incongruent − congruent.

| Pairwise Comparisons — Mov_length |  |  |  |  |  |  |  |  |
| Contrast | Goal | Distance | Estimate | SE | z | p | Lower 95% CI | Upper 95% CI |
|---|---|---|---|---|---|---|---|---|
| Left - Right | Drink | A | 0.0375 | 0.0167 | 2.243 | **0.0249** | 0.0047 | 0.0703 |
| Left - Right | Drink | B | 0.0504 | 0.0164 | 3.076 | **0.0021** | 0.0183 | 0.0825 |
| Left - Right | Drink | C | 0.0626 | 0.0172 | 3.644 | **0.0003** | 0.0289 | 0.0963 |
| Left - Right | Drink | D | 0.0142 | 0.0158 | 0.896 | 0.3704 | -0.0168 | 0.0452 |
| Left - Right | Put_Away | A | 0.0428 | 0.0162 | 2.647 | **0.0081** | 0.0111 | 0.0745 |
| Left - Right | Put_Away | B | 0.0033 | 0.0163 | 0.202 | 0.8397 | -0.0287 | 0.0353 |
| Left - Right | Put_Away | C | 0.0188 | 0.0164 | 1.144 | 0.2525 | -0.0134 | 0.0510 |
| Left - Right | Put_Away | D | -0.0167 | 0.0162 | -1.031 | 0.3026 | -0.0485 | 0.0151 |

### Absolute time to peak velocity

**Figure 2S.**
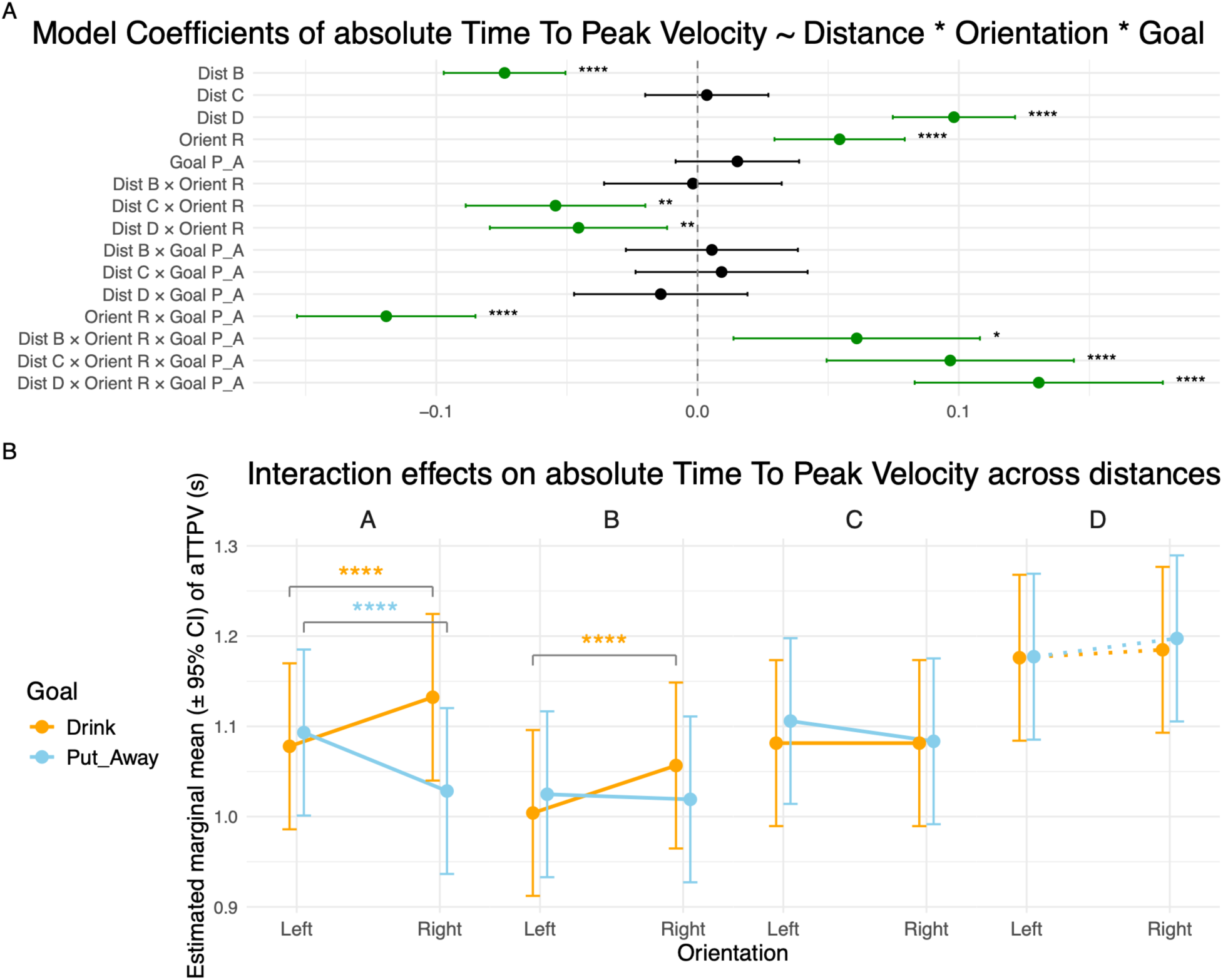
Linear Mixed Model results for Absolute Time to Peak Velocity. **A)** Forest plot of the LMM coefficients for Absolute Time to Peak Velocity, modelled as a function of Distance, Orientation, and Goal, including all interaction terms. Each row represents one predictor or interaction term, with the point estimate and 95% confidence interval displayed horizontally. Statistically significant coefficients are shown in green; the non-significant coefficient is shown in black. Asterisks denote the level of significance (* p ≤ 0.05, ** p ≤ 0.01, *** p ≤ 0.001, **** p ≤ 0.0001). **B)** Estimated marginal means (± 95% CI) of Absolute Time to Peak Velocity as a function of Orientation (Left, Right) at each distance (A, B, C, D), separately for each Goal (Drink: orange; Put Away: sky blue). Brackets with asterisks indicate planned pairwise comparisons between Left and Right orientations within each Distance × Goal combination, with the same significance coding as in panel A. Distance D is shown with dashed lines as it corresponds to a movement directed toward the right corner of the table rather than toward the mug. Note: the **congruent** combinations of Goal and Orientation are represented by Drink-Left and Put Away-Right, the **incongruent** combinations of Goal and Orientation are represented by Drink-Right and Put Away-Left.

**Table 4S.** Linear Mixed Model coefficients for Absolute Time to Peak Velocity. Fixed-effect estimates from the LMM with Absolute Time to Peak Velocity as the dependent variable, modelled as a function of Distance, Orientation, Goal, and all interaction terms. For each predictor, the table reports the coefficient estimate, standard error (SE), degrees of freedom (df), t-value, p-value, and the lower and upper bounds of the 95% confidence interval (CI). Subject ID was included as a random effect. The model intercept corresponds to the combination of Distance = A, Orientation = Left, and Goal = Drink. Distance, Orientation, and Goal are abbreviated as Dist, Orient, and Goal, respectively.

| **Model — aTTPV ~ Distance × Orientation × Goal** |  |  |  |  |  |  |  |
|---|---|---|---|---|---|---|---|
| **Fixed Effect** | **Estimate** | **SE** | **df** | **t** | **p** | **Lower 95% CI** | **Upper 95% CI** |
| (Intercept) | 1.0779 | 0.0469 | 14,9 | 22.969 | **4.73e-13** | 0.9778 | 1.1780 |
| Dist B | -0.0739 | 0.0119 | 27051 | -6.215 | **5.21e-10** | -0.0972 | -0.0506 |
| Dist C | 0.0035 | 0.0120 | 27051 | 0.294 | 0.7685 | -0.0200 | 0.0271 |
| Dist D | 0.0981 | 0.0119 | 27051 | 8.217 | **2.17e-16** | 0.0747 | 0.1215 |
| Orient R | 0.0544 | 0.0127 | 27051 | 4.274 | **1.93e-05** | 0.0294 | 0.0793 |
| Goal P_A | 0.0152 | 0.0121 | 27051 | 1.264 | 0.2064 | -0.0084 | 0.0389 |
| Dist B × Orient R | -0.0018 | 0.0173 | 27051 | -0.103 | 0.9181 | -0.0358 | 0.0322 |
| Dist C × Orient R | -0.0544 | 0.0175 | 27051 | -3.100 | **0.0019** | -0.0887 | -0.0200 |
| Dist D × Orient R | -0.0456 | 0.0173 | 27051 | -2.633 | **0.0085** | -0.0795 | -0.0116 |
| Dist B × Goal P_A | 0.0055 | 0.0168 | 27051 | 0.327 | 0.7439 | -0.0274 | 0.0384 |
| Dist C × Goal P_A | 0.0092 | 0.0168 | 27051 | 0.547 | 0.5841 | -0.0237 | 0.0421 |
| Dist D × Goal P_A | -0.0141 | 0.0169 | 27051 | -0.832 | 0.4055 | -0.0473 | 0.0191 |
| Orient R × Goal P_A | -0.1192 | 0.0174 | 27051 | -6.837 | **8.23e-12** | -0.1533 | -0.0850 |
| Dist B × Orient R × Goal P_A | 0.0609 | 0.0241 | 27051 | 2.532 | **0.0114** | 0.0138 | 0.1081 |
| Dist C × Orient R × Goal P_A | 0.0967 | 0.0241 | 27051 | 4.006 | **0.0001** | 0.0494 | 0.1440 |
| Dist D × Orient R × Goal P_A | 0.1306 | 0.0242 | 27051 | 5.389 | **7.14e-08** | 0.0831 | 0.1781 |

**Table 5S.** Pairwise comparisons of Absolute Time to Peak Velocity between Left and Right orientations. Estimated marginal mean differences (Δ = Left − Right) between orientation conditions at each Distance × Goal combination (2 Goals × 4 Distances), derived from the LMM using the emmeans package. P-values are Bonferroni-corrected for multiple comparisons. Note: For the Drink goal, Left − Right corresponds to Δ = congruent − incongruent; for the Put Away goal, Left − Right corresponds to Δ = incongruent − congruent.

| Pairwise Comparisons — aTTPV |  |  |  |  |  |  |  |  |
|---|---|---|---|---|---|---|---|---|
| Contrast | Goal | Distance | Estimate | SE | z | p | Lower 95% CI | Upper 95% CI |
| Left - Right | Drink | A | -0.0544 | 0.0127 | -4.274 | **1.92e-05** | -0.0793 | -0.0294 |
| Left - Right | Drink | B | -0.0526 | 0.0118 | -4.461 | **8.15e-06** | -0.0757 | -0.0295 |
| Left - Right | Drink | C | 2.88e-06 | 0.0121 | 2.39e-04 | 0.9998 | -0.0237 | 0.0237 |
| Left - Right | Drink | D | -0.0088 | 0.0117 | -0.749 | 0.4540 | -0.0318 | 0.0142 |
| Left - Right | Put_Away | A | 0.0648 | 0.0119 | 5.448 | **5.10e-08** | 0.0415 | 0.0881 |
| Left - Right | Put_Away | B | 0.0057 | 0.0117 | 0.486 | 0.6273 | -0.0172 | 0.0285 |
| Left - Right | Put_Away | C | 0.0224 | 0.0115 | 1.946 | 0.0517 | -0.0002 | 0.0451 |
| Left - Right | Put_Away | D | -0.0202 | 0.0120 | -1.679 | 0.0932 | -0.0438 | 0.0034 |

### Peak acceleration

**Figure 3S.**
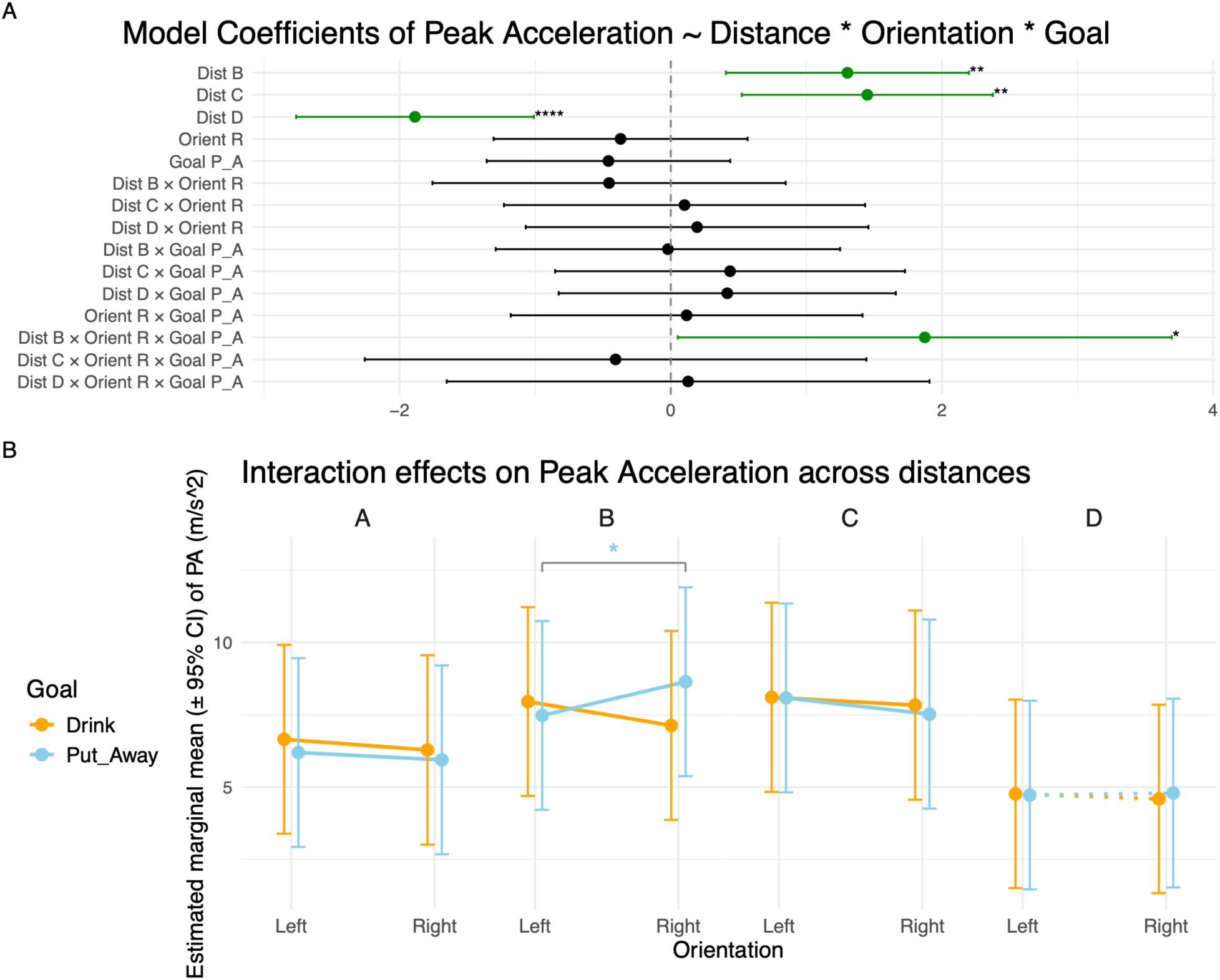
Linear Mixed Model results for Peak Acceleration. **A)** Forest plot of the LMM coefficients for Peak Acceleration, modelled as a function of Distance, Orientation, and Goal, including all interaction terms. Each row represents one predictor or interaction term, with the point estimate and 95% confidence interval displayed horizontally. Statistically significant coefficients are shown in green; the non-significant coefficient is shown in black. Asterisks denote the level of significance (* p ≤ 0.05, ** p ≤ 0.01, *** p ≤ 0.001, **** p ≤ 0.0001). **B)** Estimated marginal means (± 95% CI) of Peak Acceleration as a function of Orientation (Left, Right) at each distance (A, B, C, D), separately for each Goal (Drink: orange; Put Away: sky blue). Brackets with asterisks indicate planned pairwise comparisons between Left and Right orientations within each Distance × Goal combination, with the same significance coding as in panel A. Distance D is shown with dashed lines as it corresponds to a movement directed toward the right corner of the table rather than toward the mug. Note: the **congruent** combinations of Goal and Orientation are represented by Drink-Left and Put Away-Right, the **incongruent** combinations of Goal and Orientation are represented by Drink-Right and Put Away-Left.

**Table 6S.** Linear Mixed Model coefficients for Peak Acceleration. Fixed-effect estimates from the LMM with Peak Acceleration as the dependent variable, modelled as a function of Distance, Orientation, Goal, and all interaction terms. For each predictor, the table reports the coefficient estimate, standard error (SE), degrees of freedom (df), t-value, p-value, and the lower and upper bounds of the 95% confidence interval (CI). Subject ID was included as a random effect. The model intercept corresponds to the combination of Distance = A, Orientation = Left, and Goal = Drink. Distance, Orientation, and Goal are abbreviated as Dist, Orient, and Goal, respectively.

| **Model — PA ~ Distance × Orientation × Goal** |  |  |  |  |  |  |  |
| **Fixed Effect** | **Estimate** | **SE** | **df** | **t** | **p** | **Lower 95% CI** | **Upper 95% CI** |
|---|---|---|---|---|---|---|---|
| (Intercept) | 6.6535 | 1.6654 | 15 | 3.995 | **0.0012** | 3.1036 | 10.2033 |
| Dist B | 1.3026 | 0.4572 | 26371 | 2.849 | **0.0044** | 0.4064 | 2.1987 |
| Dist C | 1.4493 | 0.4724 | 26371 | 3.068 | **0.0022** | 0.5234 | 2.3753 |
| Dist D | -1.8865 | 0.4472 | 26371 | -4.218 | **2.47e-05** | -2.7630 | -1.0099 |
| Orient R | -0.3701 | 0.4778 | 26371 | -0.775 | 0.4385 | -1.3065 | 0.5663 |
| Goal P_A | -0.4596 | 0.4582 | 26371 | -1.003 | 0.3159 | -1.3576 | 0.4385 |
| Dist B × Orient R | -0.4552 | 0.6646 | 26371 | -0.685 | 0.4933 | -1.7578 | 0.8473 |
| Dist C × Orient R | 0.1014 | 0.6796 | 26371 | 0.149 | 0.8814 | -1.2306 | 1.4333 |
| Dist D × Orient R | 0.1941 | 0.6447 | 26371 | 0.301 | 0.7634 | -1.0696 | 1.4578 |
| Dist B × Goal P_A | -0.0212 | 0.6477 | 26371 | -0.033 | 0.9739 | -1.2907 | 1.2484 |
| Dist C × Goal P_A | 0.4370 | 0.6582 | 26371 | 0.664 | 0.5067 | -0.8531 | 1.7271 |
| Dist D × Goal P_A | 0.4164 | 0.6343 | 26371 | 0.656 | 0.5115 | -0.8269 | 1.6597 |
| Orient R × Goal P_A | 0.1167 | 0.6617 | 26371 | 0.176 | 0.8600 | -1.1803 | 1.4138 |
| Dist B × Orient R × Goal P_A | 1.8729 | 0.9292 | 26371 | 2.016 | **0.0439** | 0.0516 | 3.6943 |
| Dist C × Orient R × Goal P_A | -0.4070 | 0.9436 | 26371 | -0.431 | 0.6662 | -2.2566 | 1.4425 |
| Dist D × Orient R × Goal P_A | 0.1278 | 0.9081 | 26371 | 0.141 | 0.8881 | -1.6522 | 1.9078 |

**Table 7S.** Pairwise comparisons of Peak Acceleration between Left and Right orientations. Estimated marginal mean differences (Δ = Left − Right) between orientation conditions at each Distance × Goal combination (2 Goals × 4 Distances), derived from the LMM using the emmeans package. P-values are Bonferroni-corrected for multiple comparisons. Note: For the Drink goal, Left − Right corresponds to Δ = congruent − incongruent; for the Put Away goal, Left − Right corresponds to Δ = incongruent − congruent.

| **Pairwise Comparisons — PA** |  |  |  |  |  |  |  |  |
| **Contrast** | **Goal** | **Distance** | **Estimate** | **SE** | **z** | **p** | **Lower 95% CI** | **Upper 95% CI** |
|---|---|---|---|---|---|---|---|---|
| Left - Right | Drink | A | 0.3701 | 0.4778 | 0.775 | 0.4385 | -0.5663 | 1.3065 |
| Left - Right | Drink | B | 0.8253 | 0.4619 | 1.787 | 0.0739 | -0.0799 | 1.7306 |
| Left - Right | Drink | C | 0.2687 | 0.4835 | 0.556 | 0.5783 | -0.6789 | 1.2163 |
| Left - Right | Drink | D | 0.1760 | 0.4330 | 0.406 | 0.6844 | -0.6727 | 1.0248 |
| Left - Right | Put_Away | A | 0.2534 | 0.4576 | 0.554 | 0.5798 | -0.6435 | 1.1503 |
| Left - Right | Put_Away | B | -1.1643 | 0.4599 | -2.531 | **0.0114** | -2.0658 | -0.2629 |
| Left - Right | Put_Away | C | 0.5590 | 0.4673 | 1.196 | 0.2315 | -0.3568 | 1.4749 |
| Left - Right | Put_Away | D | -0.0685 | 0.4460 | -0.154 | 0.8779 | -0.9426 | 0.8056 |

### Absolute time to peak acceleration

**Figure 4S.**
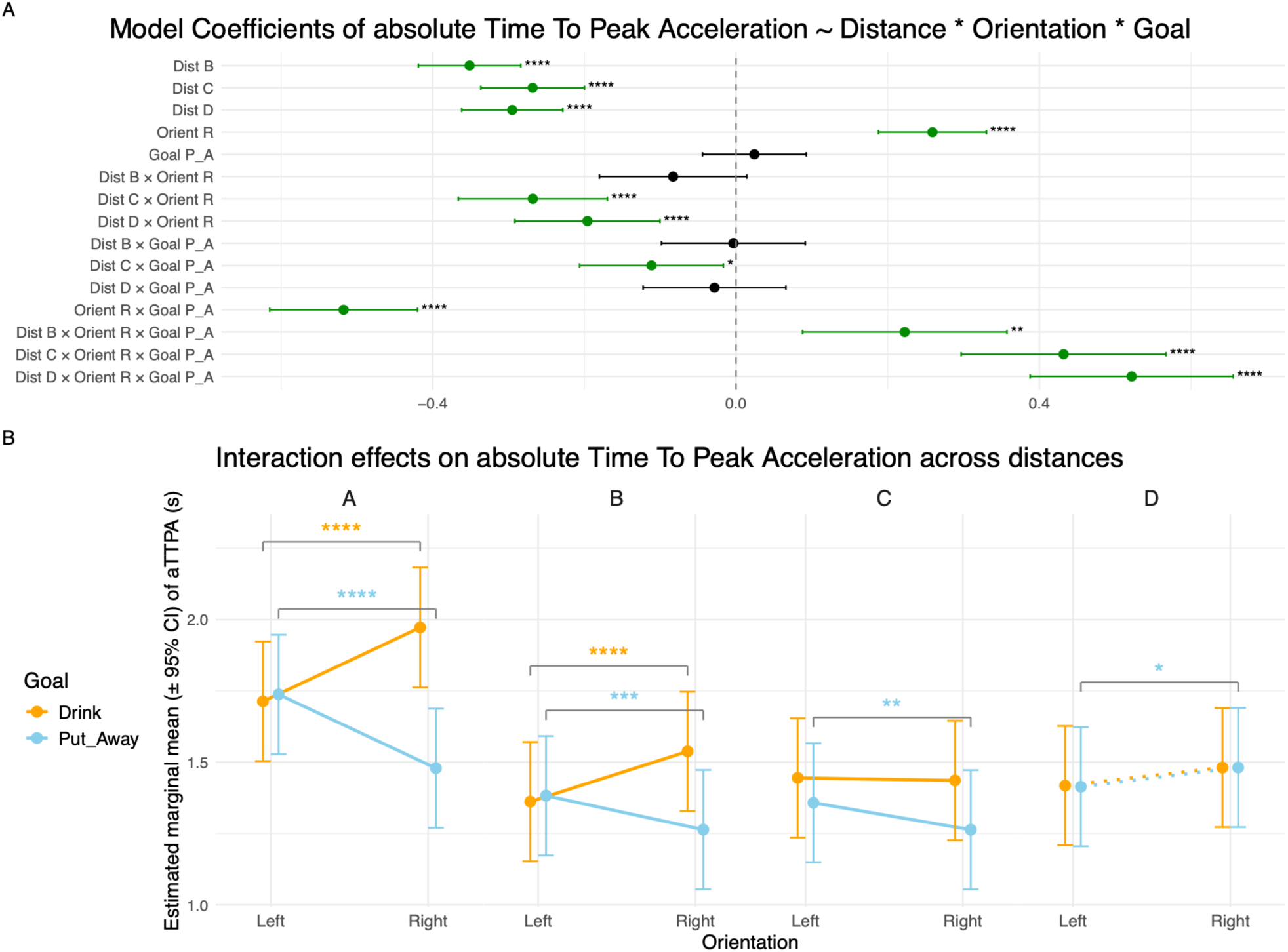
Linear Mixed Model results for Absolute Time to Peak Acceleration. **A)** Forest plot of the LMM coefficients for Absolute Time to Peak Acceleration, modelled as a function of Distance, Orientation, and Goal, including all interaction terms. Each row represents one predictor or interaction term, with the point estimate and 95% confidence interval displayed horizontally. Statistically significant coefficients are shown in green; the non-significant coefficient is shown in black. Asterisks denote the level of significance (* p ≤ 0.05, ** p ≤ 0.01, *** p ≤ 0.001, **** p ≤ 0.0001). **B)** Estimated marginal means (± 95% CI) of Absolute Time to Peak Acceleration as a function of Orientation (Left, Right) at each distance (A, B, C, D), separately for each Goal (Drink: orange; Put Away: sky blue). Brackets with asterisks indicate planned pairwise comparisons between Left and Right orientations within each Distance × Goal combination, with the same significance coding as in panel A. Distance D is shown with dashed lines as it corresponds to a movement directed toward the right corner of the table rather than toward the mug. Note: the **congruent** combinations of Goal and Orientation are represented by Drink-Left and Put Away-Right, the **incongruent** combinations of Goal and Orientation are represented by Drink-Right and Put Away-Left.

**Table 8S.** Linear Mixed Model coefficients for Absolute Time to Peak Acceleration. Fixed-effect estimates from the LMM with Absolute Time to Peak Acceleration as the dependent variable, modelled as a function of Distance, Orientation, Goal, and all interaction terms. For each predictor, the table reports the coefficient estimate, standard error (SE), degrees of freedom (df), t-value, p-value, and the lower and upper bounds of the 95% confidence interval (CI). Subject ID was included as a random effect. The model intercept corresponds to the combination of Distance = A, Orientation = Left, and Goal = Drink. Distance, Orientation, and Goal are abbreviated as Dist, Orient, and Goal, respectively.

| **Model — aTTPA ~ Distance × Orientation × Goal** |  |  |  |  |  |  |  |
| **Fixed Effect** | **Estimate** | **SE** | **df** | **t** | **p** | **Lower 95% CI** | **Upper 95% CI** |
|---|---|---|---|---|---|---|---|
| (Intercept) | 1.7132 | 0.1069 | 15,6 | 16.024 | **4.32e-11** | 1.4861 | 1.9404 |
| Dist B | -0.3514 | 0.0345 | 26847 | -10.196 | **2.28e-24** | -0.4189 | -0.2838 |
| Dist C | -0.2682 | 0.0349 | 26847 | -7.684 | **1.60e-14** | -0.3366 | -0.1998 |
| Dist D | -0.2951 | 0.0340 | 26847 | -8.676 | **4.31e-18** | -0.3618 | -0.2284 |
| Orient R | 0.2591 | 0.0363 | 26847 | 7.131 | **1.02e-12** | 0.1878 | 0.3303 |
| Goal P_A | 0.0242 | 0.0349 | 26847 | 0.695 | 0.4871 | -0.0441 | 0.0926 |
| Dist B × Orient R | -0.0829 | 0.0496 | 26847 | -1.673 | 0.0944 | -0.1801 | 0.0143 |
| Dist C × Orient R | -0.2679 | 0.0502 | 26847 | -5.340 | **9.39e-08** | -0.3663 | -0.1696 |
| Dist D × Orient R | -0.1959 | 0.0488 | 26847 | -4.013 | **0.0001** | -0.2916 | -0.1002 |
| Dist B × Goal P_A | -0.0034 | 0.0484 | 26847 | -0.071 | 0.9435 | -0.0983 | 0.0914 |
| Dist C × Goal P_A | -0.1114 | 0.0484 | 26847 | -2.303 | **0.0213** | -0.2063 | -0.0166 |
| Dist D × Goal P_A | -0.0283 | 0.0480 | 26847 | -0.590 | 0.5551 | -0.1225 | 0.0658 |
| Orient R × Goal P_A | -0.5176 | 0.0497 | 26847 | -10.411 | **2.47e-25** | -0.6150 | -0.4201 |
| Dist B × Orient R × Goal P_A | 0.2225 | 0.0688 | 26847 | 3.237 | **0.0012** | 0.0878 | 0.3573 |
| Dist C × Orient R × Goal P_A | 0.4320 | 0.0689 | 26848 | 6.268 | **3.71e-10** | 0.2969 | 0.5671 |
| Dist D × Orient R × Goal P_A | 0.5218 | 0.0683 | 26848 | 7.639 | **2.27e-14** | 0.3879 | 0.6557 |

**Table 9S.** Pairwise comparisons of Absolute Time to Peak Acceleration between Left and Right orientations. Estimated marginal mean differences (Δ = Left − Right) between orientation conditions at each Distance × Goal combination (2 Goals × 4 Distances), derived from the LMM using the emmeans package. P-values are Bonferroni-corrected for multiple comparisons. Note: For the Drink goal, Left − Right corresponds to Δ = congruent − incongruent; for the Put Away goal, Left − Right corresponds to Δ = incongruent − congruent.

| **Pairwise Comparisons — aTTPA** |  |  |  |  |  |  |  |  |
| **Contrast** | **Goal** | **Distance** | **Estimate** | **SE** | **z** | **p** | **Lower 95% CI** | **Upper 95% CI** |
|---|---|---|---|---|---|---|---|---|
| Left - Right | Drink | A | -0.2591 | 0.0363 | -7.131 | **9.96e-13** | -0.3302 | -0.1879 |
| Left - Right | Drink | B | -0.1761 | 0.0337 | -5.219 | **1.80e-07** | -0.2422 | -0.1100 |
| Left - Right | Drink | C | 0.0089 | 0.0346 | 0.257 | 0.7975 | -0.0590 | 0.0767 |
| Left - Right | Drink | D | -0.0631 | 0.0326 | -1.936 | 0.0529 | -0.1270 | 0.0008 |
| Left - Right | Put_Away | A | 0.2585 | 0.0339 | 7.623 | **2.48e-14** | 0.1921 | 0.3250 |
| Left - Right | Put_Away | B | 0.1189 | 0.0334 | 3.566 | **0.0004** | 0.0536 | 0.1843 |
| Left - Right | Put_Away | C | 0.0945 | 0.0328 | 2.881 | **0.0040** | 0.0302 | 0.1587 |
| Left - Right | Put_Away | D | -0.0673 | 0.0335 | -2.007 | **0.0447** | -0.1331 | -0.0016 |

